# AMPK Modulates Associative Learning via Neuronal Mitochondrial Fusion in *C. elegans*

**DOI:** 10.1101/2020.11.05.370031

**Authors:** Caroline C. Escoubas, Vanessa Laversenne, Emina Tabakovic, Heather J. Weir, Nicole Clark, William B. Mair

## Abstract

Loss of metabolic homeostasis is one of the hallmarks of the aging process that might contribute to pathogenesis by creating a permissive landscape over which neurodegenerative diseases can take hold. AMPK, a conserved energy sensor, extends lifespan and is protective in some neurodegenerative models. AMPK regulates mitochondrial homeostasis and morphology, however, whether mitochondrial regulation causally links AMPK to protection against loss of neuronal function with aging and diseases remains unclear. Here we use an associative learning protocol in *C. elegans* as a readout of neuronal function and show that AMPK activation enhances associative learning and prevents age-related loss of learning capacity. AMPK promotes neuronal mitochondrial fusion and mitochondrial fragmentation via *fzo-1* deletion blocks AMPK’s effects on associative learning. Restoring mitochondrial fusion capacity specifically in the neurons rescued learning capacity downstream of AMPK. Finally, AMPK activation rescues neuronal Aβ^1-42^ induced loss of associative learning. Overall, our results suggest that targeting neuronal metabolic flexibility may be a viable therapeutic option to restore neuronal function in the context of aging and neurodegenerative diseases.

## INTRODUCTION

The dramatic increase in life expectancy during the 20^th^ century was accompanied by a resultant epidemic of age-related pathologies including neurodegenerative diseases, the most prevalent of them being Alzheimer’s Disease (AD). Around 5 million individuals currently suffer from AD in the U.S., and this number is predicted to rise above 13 million by 2050(1) (www.alz.org). The available FDA-approved therapeutics are symptomatic treatments(2–4), leaving a growing need for preventive or curative drugs. The amyloid cascade hypothesis, which states that amyloid-β (Aβ) peptide deposition in the brain is a crucial event in AD pathology(5–7), is being increasingly challenged as accumulating evidence suggest that targeting the protein aggregation aspect of the disease alone is not sufficient to alleviate pathogenesis. Indeed, several Phase III clinical trials targeting Aβ and APP (Amyloid Precursor Protein) processing have failed(4). Moreover, studies show a poor association between Aβ accumulation and cognitive function(8,9), which suggests secondary factors impact the effect of Aβ on disease progression. Taken together, these observations call for a re-evaluation of our current strategy towards AD and a need to define novel risk factors.

Recent epidemiological studies have suggested a strong association between metabolic dysfunction and neurodegeneration(10)(11). Indeed, recent data show that, beyond type II diabetes, obese patients with metabolic dysfunction have increased risk of AD(10), while dietary restriction maintains metabolic homeostasis and is neuroprotective (12)(13). Loss of metabolic homeostasis is one of the hallmarks of the aging process that might contribute to disease progression by creating a permissive landscape over which neurodegenerative diseases such as AD can take hold. Our central hypothesis is therefore that targeting metabolic pathways disrupted by AD pathophysiology may be neuro-protective. Our goal is to first identify metabolic pathways contributing to age-related cognitive decline and then assess whether modulating them might have beneficial effects in an AD animal model.

AMP activated protein kinase (AMPK) is a conserved energy sensor that is activated in a low energy state when the AMP:ATP ratio increases(14)(15). Constitutive activation of AMPK extends lifespan in *C. elegans* and *Drosophila* (14,16,17) and pharmacological activation of AMPK via metformin increases healthspan in mice(14,18). Studies show a general decline in AMPK activation capacity with age under stress conditions in several tissues including the brain(19). In addition, deregulation of AMPK has been implicated in neurodegenerative diseases such as Alzheimer’s Disease, Huntington’s Disease (HD) or Parkinson’s Disease (PD)(20–26). However, the role of AMPK as a friend of foe in AD remains controversial in mammalian models. Pharmacological activation of AMPK can reduce the Aβ load and improve behavioral outcome in mouse disease models(20,27), or on the contrary, can contribute to disease pathogenesis through tau phosphorylation(21,22). These discrepancies between the behavioral and the cellular effects of AMPK in neurons highlight the importance of understanding the mechanisms of action of AMPK in AD, and to further characterize which of the many downstream processes modulated by AMPK might mediate its behavioral benefits.

AMPK is well known as an inducer of mitochondrial biogenesis, but also has intriguing emerging roles in the regulation of both metabolic flexibility and mitochondrial dynamics(28,29). Recently AMPK was also shown to slow aging by regulating mitochondrial morphology (fusion and fission), mitochondrial homeostasis, and functional coordination with peroxisomes in *C. elegans*(30). However, whether mitochondrial regulation causally links AMPK to protection against pathological outcomes of neurodegenerative disease such as AD is unknown.

A key feature of AD is loss of cognitive function and specifically memory. Here we use an associative learning protocol in the genetically tractable organism *C. elegans* as a readout of neuronal function with age and Aβ^1-42^ accumulation to define whether AMPK activation might alleviate memory loss with age and AD. We show that age-onset decline in learning and memory capacity can be halted by genetic and pharmacological activation of AMPK. We find that constitutive activation of AMPK enhances learning ability in wild type nematodes by promoting mitochondrial fusion. AMPK enhances learning and memory by promoting mitochondrial fusion in neurons; induction of fragmented mitochondrial network in neurons blocks AMPK’s beneficial effects on associative learning while restoring mitochondrial fusion rescues it. Mirroring human disease outcomes in AD, nematodes expressing the toxic amyloid peptide Aβ^1-42^ in neurons display impaired learning ability, which can be rescued by both genetic and pharmacological activation of AMPK. Our results suggest that targeting neuronal metabolic flexibility may be a viable therapeutic option to restore neuronal function in the context of neurodegenerative diseases.

## RESULTS

### Aging Suppresses Associative Learning in *C. elegans*

In order to assay neuronal function in *C. elegans* and more specifically behavioral plasticity, we used a negative associative learning assay protocol(31). This protocol is based on the simple idea that nematodes are able to modulate their chemotaxis to a neutral element in the media (NaCl) after it has been paired with a deleterious environmental cue (starvation). In this assay, pairing of NaCl in the media with the negative stimulus starvation can induce a decrease in chemotaxis towards NaCl and this behavioral change requires the simultaneous presentation of both NaCl and starvation (Fig1A). Quantification of this behavioral change reflects the neuronal function of the nematodes and can be compared to a primitive associative learning behavior(32). We conditioned *C. elegans* for 4 hours on agar plates in absence of food and in presence of NaCl. After the conditioning period, animals were transferred on chemotaxis plates, which contained a concentration gradient of NaCl across the plate. After one hour of chemotaxis, distribution of animals across the NaCl gradient is calculated (the number of nematodes at high NaCl minus the number of nematodes at low NaCl divided by the total number of worms) (*see Materials and Methods*). In order to determine the validity of this assay as an associative learning protocol, we evaluated the behavior of a population of day 1 WT *C. elegans* fed or starved, in presence or in absence of NaCl during conditioning (Fig1B) and as expected, animals starved in presence of NaCl displayed the strongest aversion towards NaCl on chemotaxis plates, suggesting that nematodes are able to associate NaCl as a negative cue only after pairing it with starvation. However, we also noticed that *C. elegans* starved in absence of NaCl also showed a moderate repulsion toward NaCl compared to fed nematodes (in presence or absence of NaCl) (Fig1B). In order to control for the effect of starvation alone, we chose to use the no *E. coli*/0mM NaCl condition as our control condition. The difference between the chemotaxis response in the control condition and the induced NaCl repulsion after conditioning (Fig1B) determines the behavioral response for a given condition, here on referred to as the “Learning Index” (Fig1B). Short-term learning has been demonstrated to rely on protein translation whereas long-term memory relies on both gene transcription and protein translation(33). To validate further our protocol, we show that *C. elegans* trained in presence of a translation inhibitor, cycloheximide, display an impaired behavioral response (SupFig1A). Conversely, exposure to the transcription inhibitor actinomycin D did not impair short-term learning capacity and even significantly improved the behavioral response (SupFig1B). Using this paradigm, we show an age dependent decline in associative learning in *C. elegans* (Fig1C,D). We observed a 50% decline in associative learning in day 3 WT adults compared to day 1 WT adults (Fig1C) as well as a complete loss of ability to form associative learning in day 5 WT adult nematodes (Fig1D), validating our assay as a readout of age-related neuronal decline in *C. elegans*.

**Figure 1:**
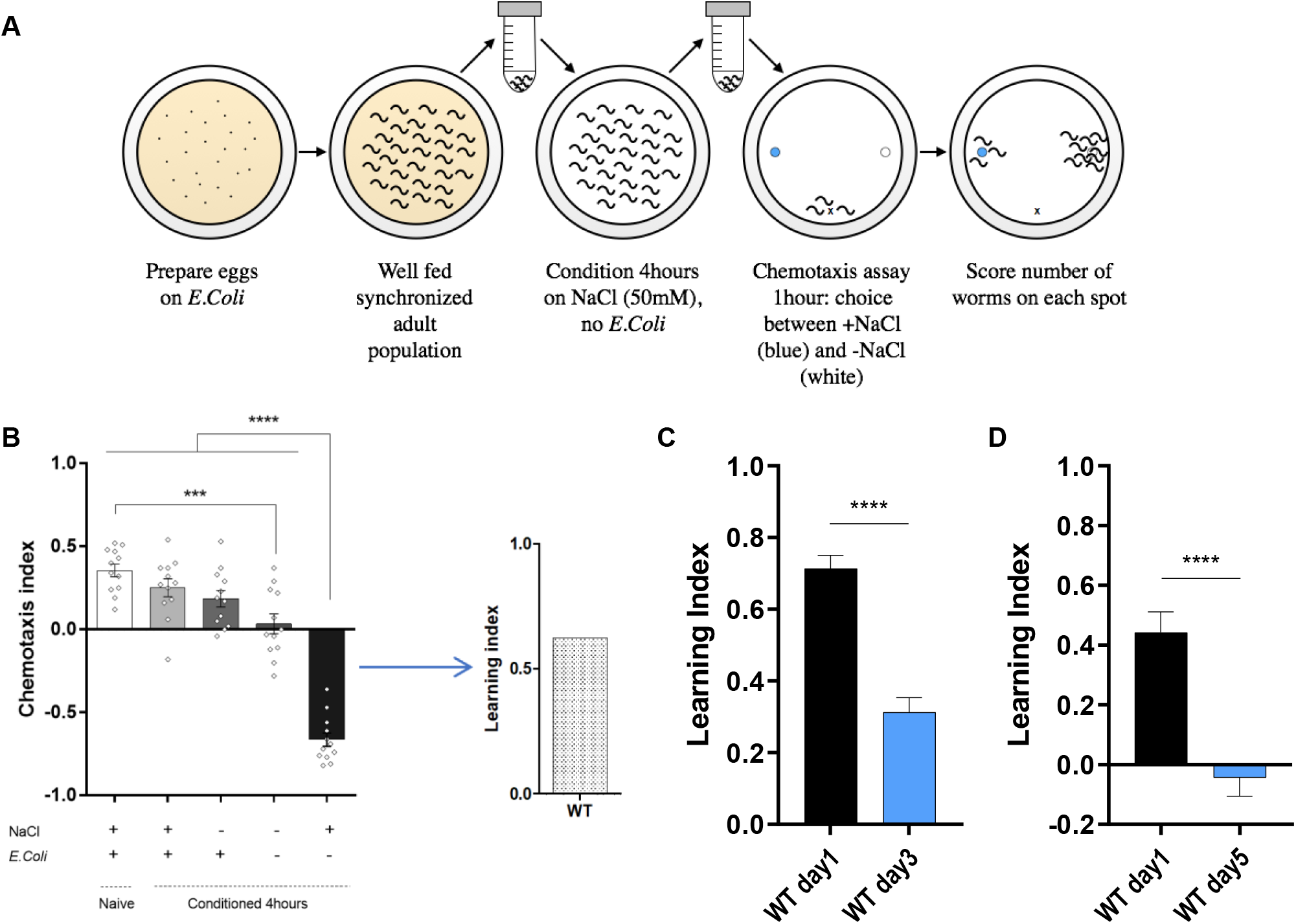
Aging suppresses associative learning in *C. elegans*. **A.** Illustration of the different steps involved in the associative learning conditioning protocol. *See Materials and Methods for detailed protocol* **B.** Chemotaxis index towards NaCl measured comparing naive (unconditioned) day 1 adult WT *C. elegans* and day 1 adult WT *C. elegans* conditioned for 4h on plates ± NaCl and ± *E. Coli*. Means naive 0.36 ± 0.04 (n=12), +NaCl +*E.Coli* 0.25 ± 0.05 (n=12), -NaCl +*E.Coli* 0.18 ± 0.05 (n=12), -NaCl -*E.Coli* 0.03 ± 0.06 (n=12), +NaCl -*E.Coli* −0.66 ± 0.04 (n=12). Comparisons naive vs -NaCl -*E.Coli p=0.0002*, +NaCl -*E.Coli* vs all other conditions *p<0.0001*. The difference between the control condition [-NaCl -*E.Coli*] and the condition [+NaCl -*E.Coli*] is referred to as the Learning Index **C.** Learning index of associative learning assay comparing WT *C. elegans* day 1 and day 3 grown on FUDR from L4 stage. Means WT day 1 adult 0.71 ± 0.04 (n=20), WT day 3 adult 0.31 ± 0.04 (n=20). Comparison WT day 1 adult vs WT day 3 adult *p<0.0001* **D.** Learning index of associative learning assay comparing WT *C. elegans* day 1 and day 5 grown on FUDR from L4 stage. Means WT day 1 adult 0.44 ± 0.07 (n=19), WT day 5 adult −0.04 ± 0.06 (n=18). Comparison WT day 1 adult vs WT day 5 adult *p<0.0001*.

### AMPK activation prevents age onset decline in associative learning

We hypothesized that the age-related decline in cognitive functions observed in humans is due to a loss of metabolic homeostasis. To begin to examine this hypothesis we asked whether modulating metabolic pathways known to extend lifespan slowed associative learning decline with age in *C. elegans*. Ubiquitous expression of a truncated catalytic subunit of AMPK (AAK-2) renders it constitutively active (referred to as CA-AMPK from here) and extends lifespan and healthspan in *C. elegans*(16,28). Interestingly, CA-AMPK *C.elegans* display an increased learning capacity at day 1 of adulthood (Fig2A). In addition to increased learning capacity after training (T=0), activation of AMPK improved memory, as assayed by the retention of the negative association after 2 hours of re-feeding on *E. coli* (T=2h) compared to WT animals (Fig2B). Next, we asked whether constitutively activating AMPK could prevent age-related loss of neuronal function. Indeed, not only did nematodes show increased learning capacity compared to WT nematodes at day 5, but more strikingly, CA-AMPK *C. elegans* did not display any age-related decline between day 1 and day 5 (Fig2C). Conversely, loss of function of the key metabolic enzyme AMPK, by mutation of its catalytic subunit *aak-2*, did not impair learning capacity in young day 1 animals (Fig2D). This result was not due to a compensation mechanism as we obtained similar results by deletion of the other isoform *aak-1* or deletion of both *aak-1* and *aak-2* isoforms (SupFig2A,B). However, by day 4 of adulthood we observed a loss of learning capacity in *aak-2^−/−^* nematodes compared to WT nematodes, although it did not quite reach statistical significance (SupFig2C *p=0.051*). The observation that *aak-2* loss of function *C. elegans* show an accelerated decline in neuronal function suggests that AMPK activity might be critical for maintaining neuronal function with age by promoting metabolic homeostasis. Finally, we asked if we could recapitulate the effects of genetic activation of AMPK with a pharmacological intervention. First, we confirmed that phenformin, the analog of the widely used drug metformin, could increase AMPK activity as measured by AAK-2 Thr-172 phosphorylation(30) and extend lifespan in an AMPK dependent manner(34) (SupFig2D). Next, we examined whether phenformin mimicked the beneficial effects of AMPK on learning. *C. elegans* grown on media supplemented with 4.5mM phenformin had enhanced learning capacity at day 1 compared to untreated nematodes (Fig2E). Pharmacological activation of AMPK therefore phenocopies its genetic activation. Taken together, these data suggest that modulation of AMPK can impact neuronal functions which normally decline with age in addition to increasing lifespan.

**Figure 2:**
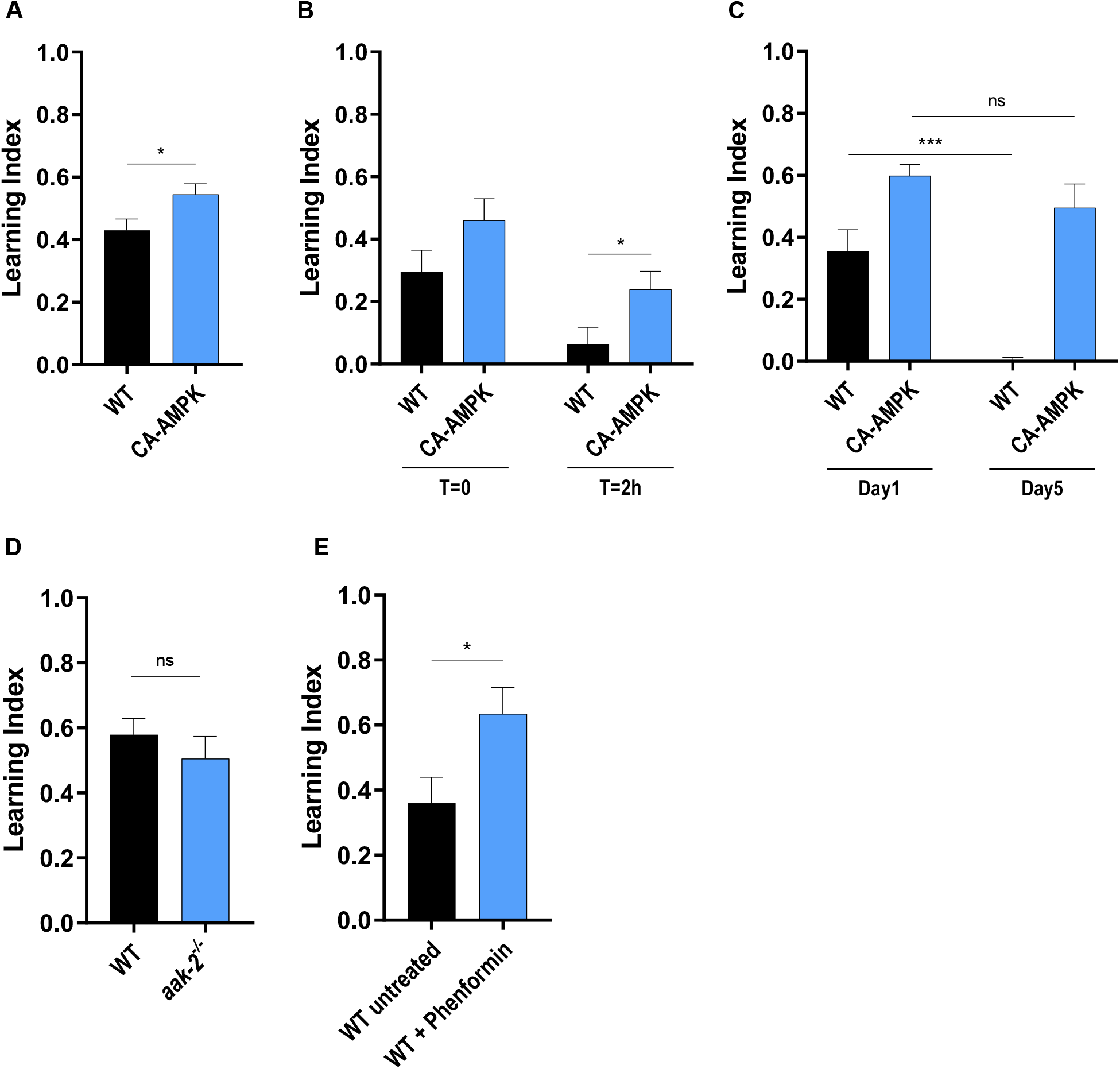
AMPK activation prevents age onset decline in associative learning. **A.** Learning index of associative learning assay comparing WT day 1 adults and CA-AMPK day 1 adults. Means WT 0.43 ± 0.04 (n=44), CA-AMPK 0.54 ± 0.03 (n=45). Comparison WT vs CA-AMPK *p=0.023* **B.** Learning index of associative learning assay comparing WT day 1 adults and CA-AMPK day 1 adults, measured immediately after conditioning (T=0h) and 2h after re-feeding on standard NGM plates seeded with OP50-1 bacteria (T=2h). Means WT T0h 0.29 ± 0.07 (n=9), CA-AMPK T0h 0.46 ± 0.07 (n=9), WT T2h 0.06 ± 0.05 (n=8), CA-AMPK T2h 0.24 ± 0.06 (n=9). Comparison WT T2h vs CA-AMPK T2h *p=0.043* **C.** Learning index of associative learning assay comparing WT vs CA-AMPK at day 1 and day 5 adulthood grown on FUDR from L4 stage. Means WT day 1 0.36 ± 0.07 (n=15), CA-AMPK day 1 0.60 ± 0.04 (n=15), WT day 5 −0.06 ± 0.07 (n=15), CA-AMPK day 5 0.50 ± 0.08 (n=15). Comparison WT day 1 vs day 5 *p=0.0003*, CA-AMPK day 1 vs day 5 *p=0.234* **D.** Learning index of associative learning assay comparing WT day 1 *C. elegans* and *aak-2^−/−^* day 1 mutants. Means WT 0.58 ± 0.05 (n=14), *aak-2^−/−^* 0.51 ± 0.07 (n=14). Comparison WT vs *aak-2^−/−^ p=0.392* **E.** Learning index of associative learning assay of WT day 1 untreated *C. elegans* vs WT day 1 *C. elegans* treated from L4 stage with 4.5mM phenformin in plate. Means WT untreated 0.36 ± 0.08 (n=13), WT phenformin 0.63 ± 0.08 (n=10). Comparison WT untreated vs WT phenformin *p=0.026*

### AMPK promotes associative learning via mitochondrial fusion

Given its key role as a homeostatic regulator of energy expenditure, AMPK modulates multiple downstream networks, ranging from transcriptional regulators, signal transduction pathways and organelle networks including mitochondrial biogenesis and dynamics(14). In *C. elegans* AMPK extends lifespan by regulating mitochondrial dynamics to maintain mitochondrial network homeostasis(28,30). Alterations in mitochondrial biology and morphology have been described in several neurodegenerative diseases, including AD. However, whether these alterations are causally implicated in age-related decline and whether manipulating mitochondrial dynamics directly can preserve neuronal function remain unknown. To explore the role of mitochondrial morphology on associative learning and its possible downstream requirement for AMPK mediated benefits on learning capacity, we first assayed worms mutated for *fzo-1*, the mammalian mitochondrial fusion factor Mfn1/2 homolog in *C. elegans*. FZO-1 regulates outer mitochondrial membrane fusion and its deletion leads to a fragmented mitochondrial network(35,36). Demonstrating a causal link between mitochondrial morphology and learning, *fzo-1* mutants have a severe learning capacity deficit at day 1 of adulthood (Fig3A). To test whether this defect was caused by deletion of the gene itself or by induction of a fragmented network we used a double deletion of *fzo-1* and *drp-1*, the *C. elegans* homolog of mammalian DRP1, which regulates mitochondrial fission(37,38) (Fig3B). Although this strain is deficient in both fusion and fission, we have previously shown that it is viable and maintains a connected mitochondrial structure(30). The double mutation can fully rescue the *fzo-1^−/−^* induced learning deficit, suggesting that *C. elegans* require a connected mitochondrial network to form associative learning (Fig3B). Induction of a “hyperfused” mitochondrial network by deletion of *drp-1* only(30) did not further enhance learning capacity in day 1 adult *C. elegans* (Fig3C). We sought to exclude the possibility that the effect observed in the mitochondrial mutant strains might be due to changes in AMPK activity levels. We performed western blot analysis of AAK-2 Thr-172 phosphorylation levels and observed a slight increase in p-AMPK which was significant in *drp-1^−/−^* and *fzo-1^−/−^* strains (SupFig3A). These results do not correlate with the behavioral phenotype of these strains and it is therefore unlikely that changes in mitochondrial network impact associative learning via modulation of AMPK activity.

**Figure 3:**
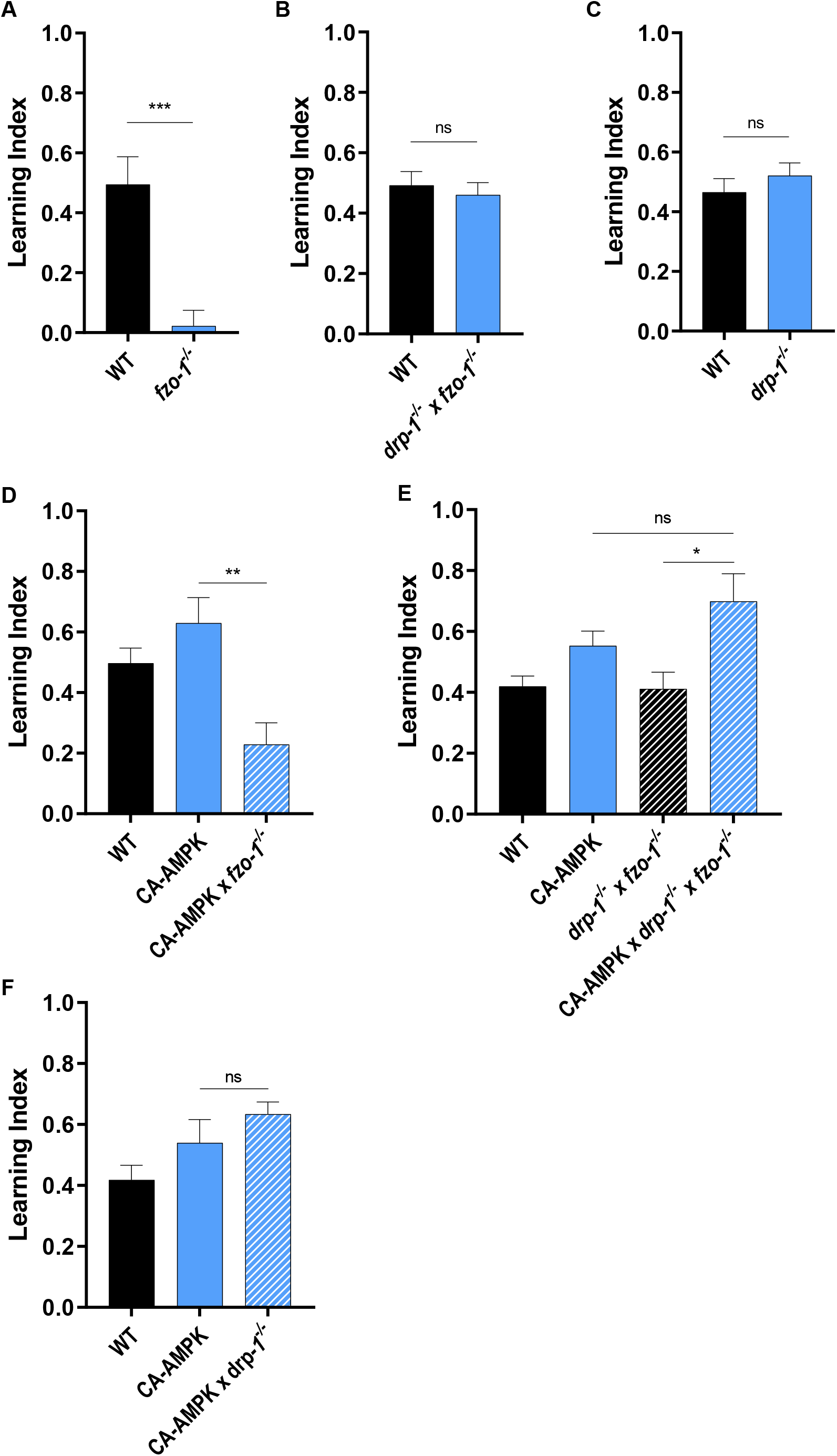
AMPK promotes associative learning via mitochondrial fusion. **A.** Learning index of associative learning assay of day 1 adult WT *C. elegans* and *fzo-1^−/−^* mutants. Means WT 0.49 ± 0.09 (n=9), *fzo-1^−/−^* 0.02 ± 0.05 (n=9). Comparison WT vs *fzo-1^−/−^ p=0.0004* **B.** Learning index of associative learning assay of day 1 adult WT *C. elegans* and *drp-1^−/−^* × *fzo-1^−/−^* mutants. Means WT 0.49 ± 0.05 (n=19), *drp-1^−/−^* × *fzo-1^−/−^* 0.46 ± 0.04 (n=20). Comparison WT vs *drp-1^−/−^* × *fzo-1^−/−^ p=0.604* **C.** Learning index of associative learning assay of day 1 adult WT *C. elegans* and *drp-1^−/−^* mutants. Means WT 0.47 ± 0.05 (n=19), *drp-1^−/−^* 0.52 ± 0.04 (n=20). Comparison WT vs *drp-1^−/−^ p=0.375* **D.** Learning index of associative learning assay of day 1 adult WT *C. elegans*, CA-AMPK *C. elegans* and CA-AMPK × *fzo-1^−/−^* mutants. Means WT 0.50 ± 0.05 (n=10), CA-AMPK 0.63 ± 0.08 (n=10), CA-AMPK x *fzo-1^−/−^* 0.23 ± 0.07 (n=10). Comparison CA-AMPK vs CA-AMPK x *fzo-1^−/−^ p=0.002* **E.** Learning index of associative learning assay of day 1 adult WT *C. elegans*, CA-AMPK *C. elegans*, *drp-1^−/−^* × *fzo-1^−/−^* mutants and CA-AMPK x *drp-1^−/−^* x *fzo-1^−/−^* mutants. Means WT 0.42 ± 0.03 (n=14), CA-AMPK 0.55 ± 0.05 (n=15), *drp-1^−/−^* x *fzo-1^−/−^* 0.41 ± 0.04 (n=14), CA-AMPK x *drp-1^−/−^* x *fzo-1^−/−^* 0.70 ± 0.09 (n=14). Comparison *drp-1^−/−^* x *fzo-1^−/−^* vs CA-AMPK x *drp-1^−/−^* x *fzo-1^−/−^ p=0.011*, CA-AMPK vs CA-AMPK x *drp-1^−/−^* x *fzo-1^−/−^ p=0.158* **F.** Learning index of associative learning assay of day 1 adult WT *C. elegans*, CA-AMPK *C. elegans* and CA-AMPK x *drp-1^−/−^* mutants. Means WT 0.42 ± 0.05 (n=14), CA-AMPK 0.54 ± 0.08 (n=14), CA-AMPK x *drp-1^−/−^* 0.63 ± 0.04 (n=14). Comparison CA-AMPK vs CA-AMPK x *drp-1^−/−^ p=0.287*.

We next asked whether mitochondrial fusion is required downstream of AMPK activation to promote associative learning. Deletion of the fusion factor *fzo-1* completely blocked CA-AMPK ability to enhance associative learning (Fig3D). This effect can be entirely rescued by simultaneous deletion of *drp-1* that restores mitochondrial connectivity, suggesting that AMPK activation requires a connected mitochondrial network to modulate neuronal function (Fig3E). Promotion of a hyper-fused mitochondrial network by deletion of *drp-1* did not show additive beneficial effects with CA-AMPK on learning capacity (Fig3F). Our observation that the double mitochondrial mutant (*fzo-1^−/−^* × *drp-1^−/−^*) displayed a WT phenotype is surprising and novel (Fig3B). Mitochondria are very dynamic organelles and work by others have suggested that their dynamic capacity is key to their function, especially in the neurons and at the synapse(39,40). Our data show that the fusion/fission process is not required in young animals to form associative learning and that the overall cellular morphology of the mitochondria is more important than its capacity to adapt through fusion or fission. This correlates with previous work from our lab(30) which demonstrated that the double mitochondrial mutant is significantly long lived. Although AMPK seems to require a balanced mitochondrial network both for longevity and promotion of neuronal functions, some differences exist; *drp-1^−/−^* blocks AMPK mediated lifespan extension(30) but did not prevent AMPK mediated associative learning (Fig3F), suggesting differential requirements for mitochondrial network state downstream of AMPK activation and potential tissue specificity of these requirements.

### Mitochondrial Fusion in Neurons is Required for Associative Learning

To distinguish between a direct requirement of mitochondrial fusion in neurons downstream of AMPK versus an indirect requirement of mitochondrial fusion mediating AMPK’s systemic beneficial effects, we created a *C. elegans* strain expressing GFP targeted to the outer mitochondrial membrane specifically in neurons (*rab-3p*:TOMM-20:GFP) in order to assess changes in neuronal mitochondrial morphology *in vivo*. In young day 1 *C. elegans*, mitochondria appeared connected and tubular and were visible both in the soma, surrounding the nucleus as well as in the dendrites projecting to the tip of the nose (Fig4A middle image). To validate our reporter, we imaged neuronal mitochondrial morphology in the mitochondrial dynamics mutant backgrounds. As expected, deletion of the outer mitochondrial membrane fusion factor *fzo-1* induces hyper-fragmentation of the network especially visible in the dendrites, as well as accumulation of aggregated signal in the soma region (Fig4A left image). Conversely, deletion of the fission factor *drp-1* causes hyperfusion of the mitochondria, which appear to form one long tubule along the whole length of the dendrite (Fig4 right image). These data demonstrate the validity and dynamic range of our reporter strain. As previously reported by other groups, neuronal mitochondrial networks become significantly more fragmented with age, as illustrated by images taken at day 1 vs. day 8 (Fig4B,C). Age induced fragmentation of the neuronal mitochondrial network correlates with the age-induced decline in learning capacity in WT animals (Fig1C,D). AMPK has previously been shown to promote mitochondrial fusion in muscles and intestine in *C. elegans* (30) and we asked whether this effect was conserved in neurons. WT worms treated from day 1 with the pharmacological AMPK activator phenformin, maintained a more tubular neuronal mitochondrial network and displayed reduced age-related fragmentation of the mitochondria at day 5 (Fig4D). To test the causal role of neuronal mitochondrial fusion in associative learning in *C. elegans*, we rescued mitochondrial dynamics specifically in the neurons by expressing *fzo-1* gene under a pan-neuronal promoter (*rab-3* promoter) in a *fzo-1* deletion background (Fig4E). Expression of neuronal FZO-1 fully rescued the learning deficit observed in fusion deficient animals (Fig4E), suggesting that mitochondrial fusion is required in neurons to promote learning capacity. Furthermore, restoring *fzo-1* expression in neurons of CA-AMPK *C. elegans* crossed with *fzo-^1−/−^* rescued its learning capacity to WT levels (Fig4F). Taken together, these results causally link the beneficial effect of AMPK on associative learning to cell autonomous modulation of neuronal mitochondrial networks in *C. elegans*.

**Figure 4:**
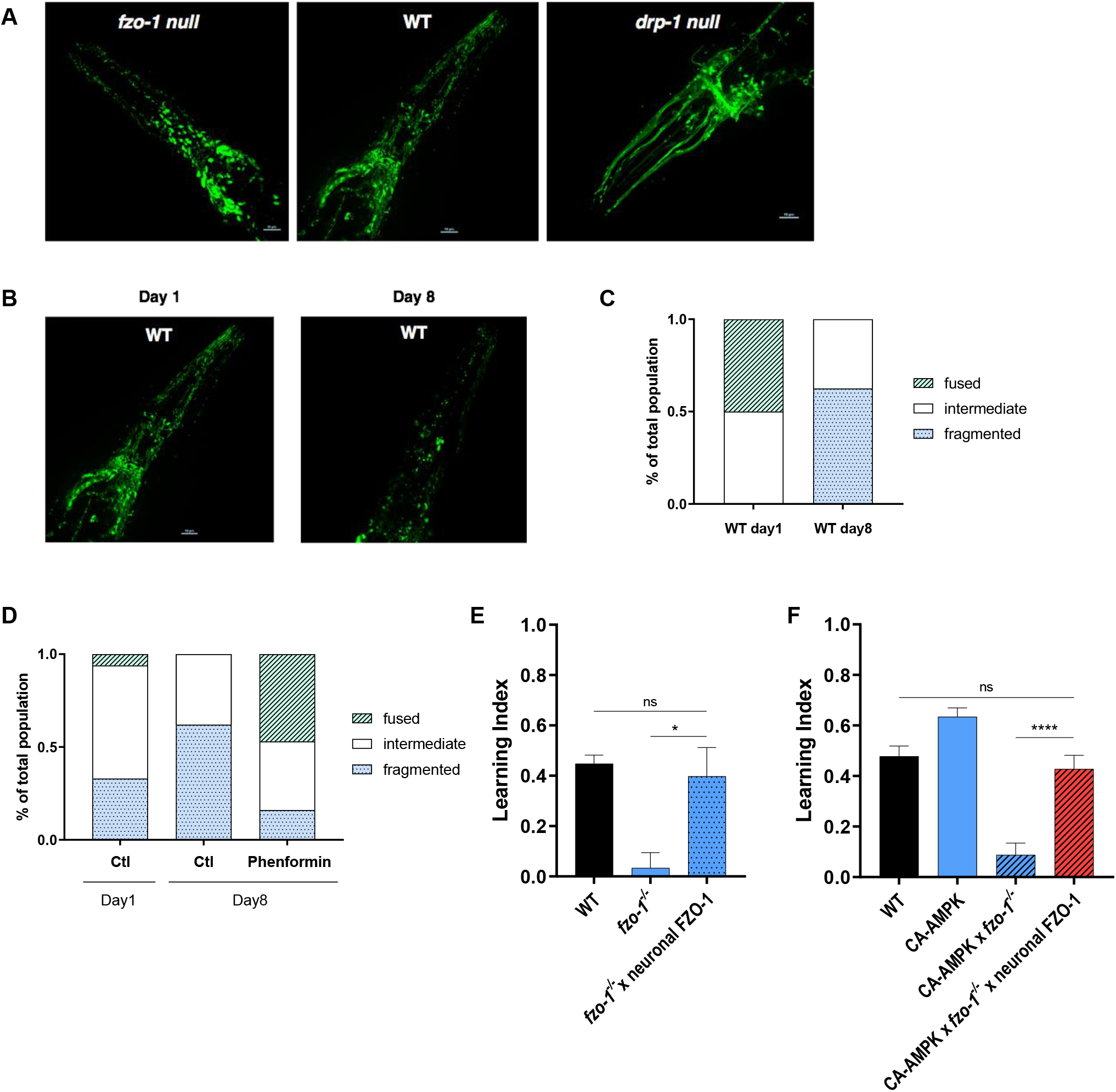
Mitochondrial fusion in neurons is required for associative learning. **A.** Representative images of the pan-neuronal mitochondrial reporter [*rab-3p*:TOMM-20:GFP] in day 1 adult *C. elegans* in WT (middle image), *fzo-1^−/−^* (left image) or *drp-1^−/−^* (right image) backgrounds. Image shows the head portion of the nematode. An average of 10 nematodes were imaged **B.** Representative images of the pan neuronal mitochondrial reporter [*rab-3p*:TOMM-20:GFP] in day 1 and day 8 adult WT *C. elegans* showing age related neuronal mitochondrial fragmentation **C.** Qualitative quantification of neuronal mitochondrial reporter [*rab-3p*:TOMM-20:GFP] in day 1 adults (n=6) and day 8 adults (n=8). Images were blinded and the mitochondrial network was scored as fused, intermediate or fragmented **D.** Representative images of the pan neuronal mitochondrial reporter [*rab-3p*:TOMM-20:GFP] in day 1 and day 8 adult WT *C. elegans* and day 8 WT *C. elegans* treated from day 1 with phenformin 4.5mM **E.** Qualitative quantification of neuronal mitochondrial reporter [*rab-3p*:TOMM-20:GFP] in day 1 adults untreated (n=18), day 8 adults untreated (n=13) and day 8 adults treated with phenformin 4.5mM from day 1 (n=19). 2 biological replicates. Images were blinded and the mitochondrial network was scored as fused, intermediate or fragmented **F.** Learning index of associative learning assay of day 1 adult WT *C. elegans*, *fzo-1^−/−^* mutants and *fzo-1^−/−^* x [rab-3p:fzo-1 cDNA]. Means WT 0.45 ± 0.03 (n=12), *fzo-1^−/−^* 0.03 ± 0.06 (n=10), *fzo-1^−/−^* x neuronal FZO-1 0.40 ± 0.11 (n=11). Comparison *fzo-1^−/−^* vs *fzo-1^−/−^* x neuronal FZO-1 *p=0.013*, WT vs *fzo-1^−/−^* x neuronal FZO-1 *p=0.668* **G.** Learning index of associative learning assay of day 1 adult WT *C. elegans*, CA-AMPK *C. elegans*, CA-AMPK x *fzo-1^−/−^* mutants and CA-AMPK x *fzo-1^−/−^* x neuronal FZO-1 mutants. Means WT 0.48 ± 0.04 (n=23), CA-AMPK 0.63 ± 0.03 (n=23), CA-AMPK x *fzo-1^−/−^* 0.09 ± 0.05 (n=20), CA-AMPK x *fzo-1^−/−^* x neuronal FZO-1 0.43 ± 0.05 (n=12). Comparisons WT vs CA-AMPK x *fzo-1^−/−^* x neuronal FZO-1 *p=0.465*, CA-AMPK x *fzo-1^−/−^* vs CA-AMPK x *fzo-1^−/−^* x neuronal FZO-1 *p<0.0001*

### Activating AMPK Restores Learning Capacity in a *C. elegans* AD model

We next aimed to translate these findings in the context of a disease model and more specifically, a model of Alzheimer’s Disease (AD). Multiple models of AD in both *C. elegans* and mouse use high expression of toxic levels of Aβ^1-42^ which result in very short lifespans and artificially severe detrimental phenotypes. To circumnavigate these concerns, we sought a model with very low expression of Aβ. We took advantage of a previous model of neurodegeneration which over-expresses the human sequence of the toxic amyloid peptide causal in Alzheimer’s disease, Aβ^1-42^, in all *C. elegans* neurons(41). This Aβ^1-42^ *C elegans* model predominantly expresses the truncation product Aβ^3-42^ which also aggregates and forms fibrillar structures(42). This strain was originally designed to be inducible, via leveraging temperature sensitive (TS) mutations in the Non-sense Mediated Decay (NMD) pathway. The Aβ^1-42^ mRNA is tagged with an extended 3’UTR that targets it to the NMD pathway for degradation. When combined with TS mutations in the NMD component *smg-1*, Aβ^1-42^ can be expressed at high levels at 25°C when SMG-1 is inactive. At low temperatures when TS SMG-1 is active, the NMD pathway is functional and the Aβ^1-42^ mRNA is targeted for degradation, leading to lower levels of Aβ^1-42^ expression (Fig5A). As we sought to have a low expression of Aβ^1-42^ we removed the TS *smg-1* allele from this line, leaving the long 3’UTR neuronal Aβ^1-42^ in a WT background. We confirmed this system by measuring expression of Aβ^1-42^ with and without RNAi knockdown of *smg-1*. As expected, when the NMD pathway is disrupted, Aβ^1-42^ expression increased (SupFig4A). However, despite the low protein expression of the amyloid transgene in the WT background with fully functional NMD, we detected the presence of Aβ in the neurons via immunohistochemistry using the 6e10 anti-Aβ antibody (low Aβ) (Fig5B).

**Figure 5:**
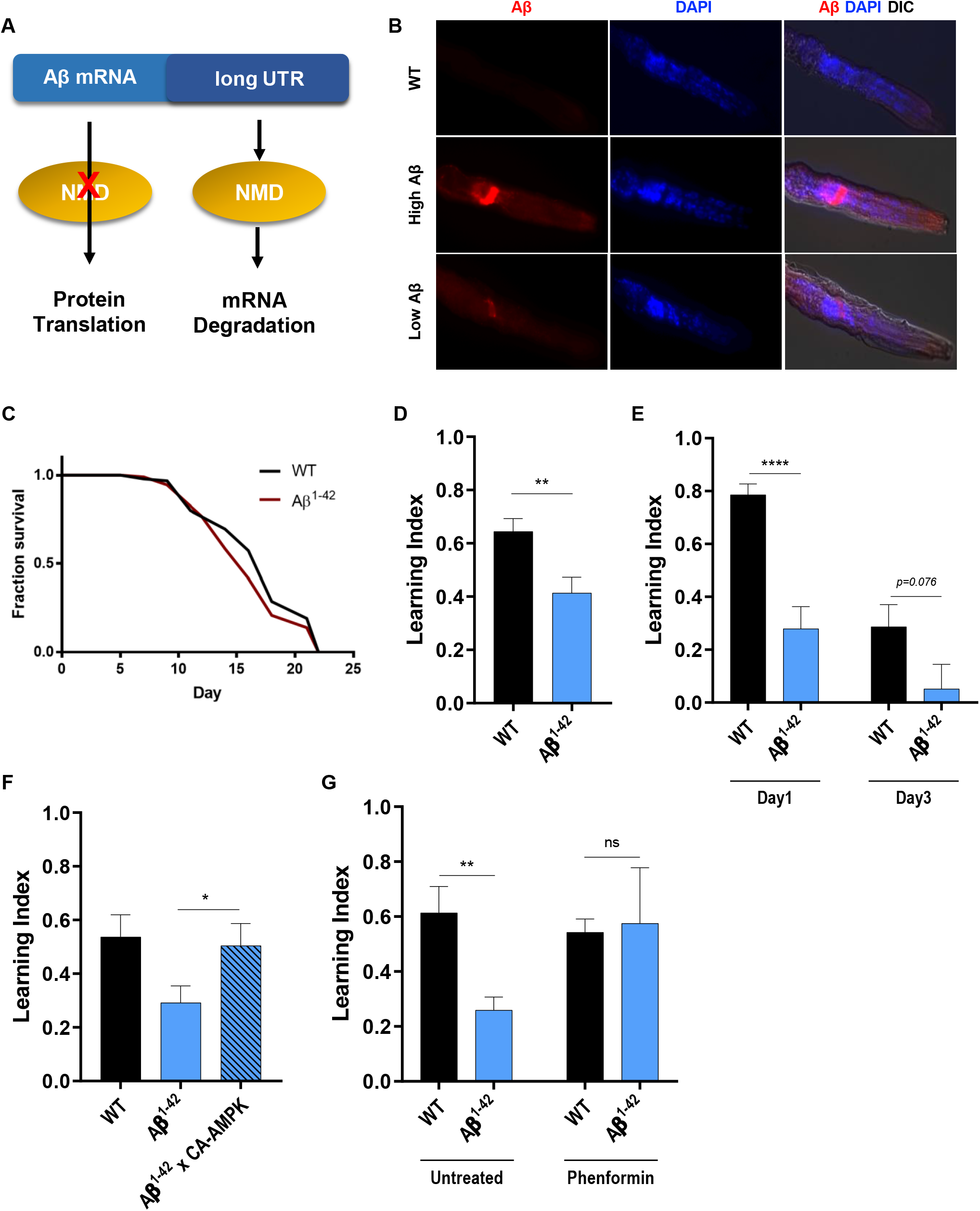
Activating AMPK restores learning capacity in a *C. elegans* AD model. **A.** Schematic representation of the Aβ^1-42^ transgene tagged with the extended 3’UTR, targeting its mRNA to the Nonsense Mediated Decay pathway **B.** Representative images of day 1 adult *C. elegans* stained with amyloid peptide antibody. Head of the nematode is shown in picture. An average of 20 nematodes were imaged. **C.** Lifespan assay of WT *C. elegans* vs Aβ^1-42^ model [Psnb-1::Abeta 1-42::3’UTRlong], *p=0.313* (log-rank Mantel-Cox test). 3 biological replicates, statistical analysis of all lifespan experiments in SupTable2. **D.** Learning index of associative learning assay in day 1 adult WT *C. elegans* and Aβ^1-42^ strain. Means WT 0.64 ± 0.05 (n=23), Aβ^1-42^ strain 0.41 ± 0.06 (n=23). Comparison WT vs Aβ^1-42^ strain *p=0.004* **E.** Learning index of associative learning assay in day 1 and day 3 adult WT *C. elegans* and Aβ^1-42^ strain. Means WT day 1 0.79 ± 0.04 (n=10), Aβ^1-42^ day 1 0.28 ± 0.08 (n=9), WT day 3 0.29 ± 0.08 (n=10), Aβ^1-42^ day 3 0.05 ± 0.09 (n=10). Comparisons WT day 1 vs Aβ^1-42^ day 1 *p<0.0001*, WT day 3 vs Aβ^1-42^ day 3 *p=0.076* **F.** Learning index of associative learning assay in day 1 adult WT *C. elegans*, Aβ^1-42^ strain and CA-AMPK x Aβ^1-42^ strain. Means WT 0.54 ± 0.08 (n=14), Aβ^1-42^ strain 0.29 ± 0.06 (n=14), CA-AMPK x Aβ^1-42^ strain 0.50 ± 0.08 (n=14). Comparison Aβ^1-42^ strain vs CA-AMPK x Aβ^1-42^ strain *p=0.049* **G.** Learning index of associative learning assay of day 1 WT and Aβ^1-42^ strain, untreated and treated with Phenformin 4.5mM from L4 stage. Means WT untreated 0.61 ± 0.10 (n=14), Aβ^1-42^ strain untreated 0.26 ± 0.05 (n=15), WT phenformin 0.54 ± 0.05 (n=14), Aβ^1-42^ strain phenformin 0.57 ± 0.20 (n=14). Comparisons WT untreated vs Aβ^1-42^ strain untreated *p=0.002*, WT phenformin vs Aβ^1-42^ strain phenformin *p=0.877*

Having developed a low Aβ^1-42^ *C. elegans* model we sought to examine its negative impact on aging itself. Unlike other published *C. elegans* AD models which display shortened lifespan, this low-level expression strain does not reduce longevity (Fig5C). Any phenotype seen in this strain are therefore likely due to a specific effect of Aβ^1-42^ peptide expression in neurons rather than overall deterioration of the nematodes’ health and accelerated aging. Using our associative learning paradigm, we demonstrated that young day 1 AD *C. elegans* display a significant impairment in their learning capacity compared to WT *C. elegans* (Fig5D). This deficit is further exacerbated by age as shown in Fig5E where, by day 3, AD worms completely lose their learning capacity, thus mimicking the age-related cognitive decline observed in AD. Strikingly, learning capacity of AD nematodes can be rescued to WT levels by CA-AMPK (Fig5F), suggesting that the detrimental effect of a proteotoxic peptide on neuronal functions can be alleviated by modulation of a key cellular energy sensor. We assessed Aβ^1-42^ mRNA levels by qPCR to verify that transgene expression was not decreased by CA-AMPK overexpression (SupFig4B). We further demonstrate that pharmacological activation of AMPK via phenformin treatment also rescues Aβ^1-42^ induced learning deficit in young *C. elegans* (Fig5G). Conversely, deletion of the catalytic subunit of AMPK, *aak-2*, seemed to further impair our AD nematodes associative learning although it did not reach statistical significance (*p value 0.055*) (SupFig4C). Having previously shown that AMPK requires fusion of the mitochondria to enhance neuronal function, we hypothesized that directly promoting mitochondrial fusion by deleting *drp-1* might have beneficial effects on our AD model. Loss of function *fzo-1* mutation did not display additional learning defects with Aβ^1-42^ expression (SupFig4D), however *drp-1* deletion failed to rescue Aβ^1-42^ induced learning defects (SupFig4E). This result suggests that either mitochondrial fusion is required to mediate AMPK beneficial effect on associative learning but not sufficient by itself to restore Aβ^1-42^-induced neuronal dysfunction.

### The Beneficial Effects of AMPK are Specific to AD neurodegenerative models

Finally, we asked whether activation of AMPK had a beneficial effect on *C. elegans* proteotoxic stress models in general or a more specific action on Aβ^1-42^ induced neuronal toxicity. We turned towards another degeneration model in *C. elegans*, a Huntington’s Disease (HD) model. HD is an autosomal dominant disease caused by an increased repeat of CAG codons in the huntingtin (Htt) gene (The Huntington’s Disease Collaborative Research Group, 1993). Similarly to AD, HD is associated with important metabolic dysfunction(43) such as ATP synthesis, mitochondrial respiration(44) and AMPK signaling impairments(24). We obtained strains expressing pan-neuronally peptides with different numbers of glutamine repeats (Q): a control strain Q0, and disease model strains Q40 and Q67(45,46). In humans, 40 CAG repeats, or more, leads to development of the disease(47) and onset of HD directly correlates with total number of repeats(48). Previous labs have demonstrated that a number of 40 CAG repeats is the threshold at which the protein shifts from being soluble in the cytoplasm to insoluble and aggregated(45,46). As expected, polyQ40 worms show some isolated punctae (SupFig5A middle image) whereas poly67 worms display complete aggregation (SupFig5A right image). Rather than looking at paralysis as a readout of neuronal dysfunction as has been done previously, we used our associative learning paradigm to assess functional impairment induced by polyQ peptide aggregation. The polyQ40 strain, which displays mild to no aggregation, shows a small but not significant decrease in learning capacity, unlike the polyQ67 strain which displayed a major loss of learning capacity (SupFig5B), similar that observed in our AD model (Fig5D). These results suggest a correlation between the extent of the polyQ aggregation and neuronal dysfunction. Genetic activation of AMPK, however, did not rescue this functional impairment (SupFig5C), suggesting that AMPK activation does not broadly reverse proteotoxic stress but rather has a beneficial effect specific to Aβ^1-42^ induced neuronal toxicity. Indeed, despite the fact that both our AD and HD model strains represent proteotoxic stress models with neuronal expression of aggregated proteins, the same intervention, AMPK activation, does not equally benefit both disease models. This observation would suggest that different cellular functions are altered downstream of Aβ^1-42^ or polyQ67 accumulation and that AMPK mediates its effect on those downstream processes. AMPK activation does not represent a universal intervention to treat various toxic aggregation disorders but seems to be more specific to the physiopathology of AD and to age-related neuronal decline. Our results nicely correlate with human and mice striatum data where AMPK was found to be abnormally activated and further contributing to oxidative stress and HTT induced neuronal cell death(24).

## DISCUSSION

Taken together, our results highlight a novel link between AMPK activation and modulation of mitochondrial fusion in the nervous system to promote behavioral plasticity. Our data suggest that AMPK activation provides beneficial effects on learning capacity in aged *C. elegans* by promoting mitochondrial fusion in the neurons. Indeed, fusion deficient worms show impaired learning, which could be rescued by restoring mitochondrial fusion specifically in the neurons. We further showed that nematodes expressing the toxic amyloid peptide Aβ^1-42^ in the neurons display impaired learning ability, which could be rescued by both genetic activation of AMPK as well as pharmacological activation of AMPK via the metformin analog, phenformin.

We chose the model system *C. elegans* for our study for the availability of genetic tools and the ease to generate transgenic animals. The transparency of the nematodes also conferred a distinct advantage as we were able to image neuronal mitochondrial morphology *in vivo* and thus to explore the link between AMPK and mitochondrial network. One major aspect which differentiates the nematode nervous system from the mammalian brain, is the fact that *C. elegans*, as a post mitotic organism, does not experience neurogenesis in the adult nematode. Adult hippocampal neurogenesis, or the production and integration of new neurons in the brain, which mainly occurs in the dentate gyrus in mammals(49), has been linked to learning and memory performance in both aversive association paradigms(50–52) and positive association paradigms(53), and is considered to have a crucial role in the hippocampal learning process. Although we cannot assess the relative contribution of adult neurogenesis to learning capacity in our system, we still observed a plasticity of the behavior in *C. elegans* after conditioning, suggesting that we are studying the intra-cellular processes involved in regulation of behavioral plasticity. *C. elegans* represents a valuable tool to study neurobiology as well as gain insight into the cellular and molecular processes disrupted in age related disorders however it cannot model all aspects of neurodegenerative disorders pathophysiology and it will be necessary to validate those findings not only in rodents but in more complex organisms such as primates or humans(54).

AMPK plays an important role in neuronal physiology in mammals, and evidence in mice suggests a beneficial role of AMPK activation in memory formation. Here we show that AMPK not only prevents age-related decline in neuronal functions but also alleviates Abeta induced loss of learning capacity. However, whether AMPK might benefit or impair cognitive outcomes in neurodegenerative diseases remains unclear. An increasing number of publications suggest a very prominent role for AMPK both for the normal function of neuronal tissue and in the context of cognitive impairment and neurodegenerative diseases(14,55–57). A cross sectional population study in Korean subjects aged >60 years old shows a significant association between the AMPK γ2 gene polymorphism (PRKAG2 −26C/T) and cognitive impairment in old age. Furthermore, AMPK is a modulator of Long Term Potentiation (LTP)(58) and old mice display a decrease in the levels of LKB1 – an AMPK upstream activator – as well as a decrease in active AMPK(59). Pointing to an important function of AMPK in synapse formation, deletion of either kinase leads to aberrant axonal and dendrite morphology whereas genetic and pharmacological (metformin, caloric restriction) activation of AMPK protects against age-related synaptic alterations in those neurons(59). AMPK proved to have an even more prominent role in the context of neurodegenerative diseases and more specifically Alzheimer’s Disease. AMPK is required downstream of two interventions known to ameliorate the pathology of Alzheimer’s Disease: leptin(20) and resveratrol(27). Inhibition of Aβ production and tau phosphorylation by leptin and extracellular Aβ accumulation by resveratrol in neuronal cultures both required AMPK and could be replicated using pharmacological activators of AMPK(20,27). Likewise, AMPK activation inhibits palmitate-induced apoptosis and tau hyperphosphorylation in SH-SY5Y cells(60). However, contradicting the potential beneficial role of AMPK in Alzheimer’s Disease, AMPK is a tau kinase, which could potentially contribute to disease pathogenesis by altering microtubule binding of tau(21). AMPK abnormally accumulates in tangle and pre-tangle bearing neurons in major tauopathies and it was shown *in vitro* that Aβ^1-42^ induced activation of AMPK via CaMKKβ (Ca2+/calmodulin-dependent protein kinase kinase β) leads to increased tau phosphorylation(21,22). These observations were confirmed *in vivo* in an AMPKα2 deficient mouse model, which displayed reduced levels of endogenous tau phosphorylation(61). Despite convincing evidence that AMPK is a tau kinase, these findings do not address the functional impact of such phosphorylation *in vivo* on cognitive functions. It is therefore difficult to assess whether this increased tau phosphorylation by AMPK contributes to the memory loss or neurodegeneration observed in Alzheimer’s Disease or instead reflects activation of a stress response. *Ma et al*. sought to study the effect of AMPK inhibition on long term potentiation (LTP) and long-term depression (LTD) in APP/PS1 transgenic mice, a process considered as the cellular basis for learning and memory formation. They showed a negative contribution of AMPK, as treatment of this murine AD model with the AMPK inhibitor compound C or genetic deletion of the AMPK catalytic subunit prevented Aβ induced LTP loss(62). This once again contrasts with the behavioral data clearly showing beneficial effects of activation of AMPK on memory formation(63,64). These discrepancies between the behavioral and the cellular effects of AMPK in neurons point to the gap in the understanding of the mechanisms of action of AMPK and the need to further characterize the downstream targets of AMPK mediating its behavioral benefits. Many studies use Aβ^1-42^ accumulation as a readout of the efficacy of the tested intervention and although Aβ peptide accumulation is a hallmark of AD, it does not always reflect the severity of the disease. We therefore used the measurement of neuronal functions, more specifically behavioral plasticity, as our main outcome and we showed a beneficial role of AMPK activation. Our findings do not exclude a potential detrimental role of AMPK activation as we could not evaluate the effect of AMPK modulation on tau pathology in our system. Several *C. elegans* models expressing pathological levels of tau in the neurons are available and it would be interesting to test whether AMPK activation can benefit those models.

Here we defined mitochondrial fusion as a required downstream mechanism for AMPK mediated beneficial effect on cognitive function in the context of aging and neurodegeneration. Mitochondrial biology is key to neuronal physiology, has been linked to cognitive disorders and is therefore a relevant target to study downstream of AMPK. However, AMPK is a master regulator of metabolism in the cell and regulates a vast array of other processes, which might contribute to the effect of AMPK modulation in the context of neurodegeneration. AMPK regulates autophagy by interacting with ULK1/2 and Beclin-1 and this catabolic response to starvation serves one purpose, the recycling of cytosolic content such as the organelles(65). Autophagy would be a relevant downstream mechanism to study here as it plays a role in physiological degradation of the amyloid peptides contained in endosomes in the brain(66). Autophagy is also dysregulated in AD as mutation of the Prenisilin 1 gene disrupts autophagy by inducing a loss of lysosomal acidification(67). Deletion of Beclin-1, a downstream target of AMPK, leads to increased Aβ accumulation in AD mice models while its overexpression alleviates the proteotoxic accumulation in cultured neurons(68,69). Inducing autophagy therefore represents a physiological way to degrade the accumulated aggregates and slow the progression of the disease. It would be interesting to determine whether induction of autophagy contributes to the beneficial behavioral effect observed with AMPK activation in our models.

Neurons are very energy dependent and require a tight regulation of calcium homeostasis for synaptic function thus making them very sensitive to modification in mitochondrial function. Given the importance of mitochondrial function and distribution in neuronal cells, it is not surprising that disruption of mitochondrial dynamics machinery leads to neuronal dysfunction and neurological diseases(39). We show here that induction of mitochondrial fragmentation leads to loss of learning capacity in young nematodes and that AMPK beneficial effects are mediated by neuronal mitochondrial fusion. Our results are consistent with previously published studies suggesting a link between dysregulation of mitochondrial function or mitochondrial dynamics and neurodegenerative disorders such as AD, HD and PD(11). In AD, multiple aspects of mitochondrial function have been shown to be altered (11); AD brains display a deficiency in TCA (tricarboxylic acid or Krebs) cycle enzymes(70), impaired PGC1α expression contributes to AD progression(71), Prenisilin 1 mutation causes mitochondrial axonal transport defects(72), Aβ peptide accumulates on and blocks mitochondrial import channels(73), increased percentage of mtDNA mutations has been described in AD brains(74) and calcium homeostasis is severely altered(75,76).

Furthermore, neuronal expression of mitochondrial dynamics machinery protein is significantly reduced in AD hippocampi(77). Mitochondrial dynamics are also altered in AD, as earlier studies done in AD neurons described modifications of mitochondrial size and number as well as an increase in mitochondria with broken cristae(74). Exposure of cultured neurons to aggregated Aβ peptide or overexpression of APP and APPsw mutant can also lead to mitochondrial fragmentation(77–79). Alteration of mitochondrial capacity to fuse or divide consequently leads to axonal transport defect and abnormal mitochondrial distribution between the soma and the dendrites(40,77–79). Interestingly, the activity of a rate limiting enzyme in the TCA cycle, α-ketoglutarate dehydrogenase, is reduced in a *C. elegans* strain expressing neuronal Aβ. This dysregulation in mitochondrial metabolic function is rescued by metformin treatment suggesting that modulation of TCA cycle intermediates could be a potential mechanism though which AMPK promotes beneficial effects(80). Although we clearly demonstrate a requirement for mitochondrial fusion downstream of AMPK specifically in the neuronal tissue, we cannot exclude a systemic contribution of AMPK activation. Even though the brain seems to be the tissue of interest to target, a systemic action of AMPK activation could be very interesting when considering translational potential as crossing the blood brain barrier in humans represents one of the main problems in developing drugs for neurological disorders. Although the AMPK activator, metformin, has been shown to cross the blood brain barrier in mice, metformin was delivered systemically in all studies to the animals and also improved metabolic outcomes, such as insulin resistance and glucose tolerance.

In conclusion, our results suggest that AMPK is a key enzyme in maintaining learning capacity with age in *C. elegans* and further rescues Aβ^1-42^ induced detrimental effect on this functional outcome. Interventions targeting metabolic dysfunction and more specifically AMPK downstream mechanisms such as mitochondrial dynamics may therefore be beneficial to alleviate AD pathogenesis and functional impairment.

---

**Supplemental Figure 1:**
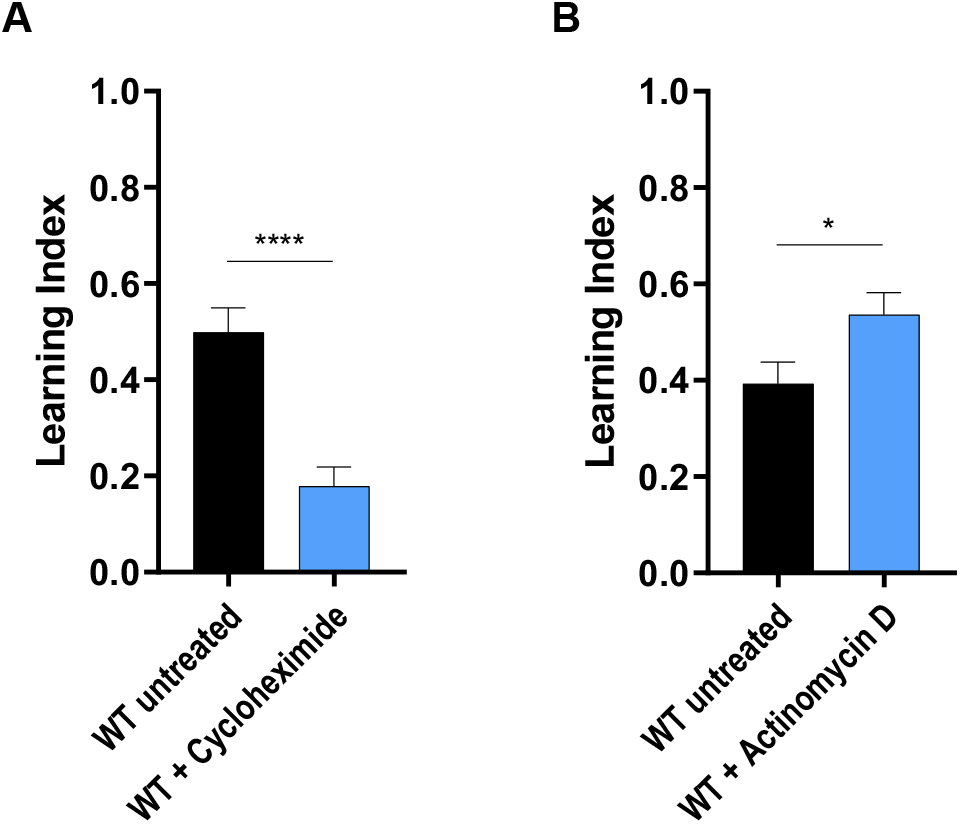
Short term associative learning in *C. elegans* requires translation. **A.** Learning index of associative learning assay comparing day 1 adult WT *C. elegans* untreated to day 1 adult WT *C. elegans* treated for 4h on plates containing cycloheximide (800μg/mL). Means untreated 0.50 ± 0.05 (n=25), cycloheximide treated 0.18 ± 0.04 (n=24). Comparison untreated vs cycloheximide treated *p<0.0001* **B.** Learning index of associative learning assay comparing day 1 adult WT *C. elegans* untreated to day 1 adult WT *C. elegans* treated for 4h on plates containing actinomycin D. Means untreated 0.39 ± 0.05 (n=21), actinomycin D treated 0.54 ± 0.05 (n=21). Comparison untreated vs actinomycin D treated *p=0.032*.

**Supplemental Figure 2:**
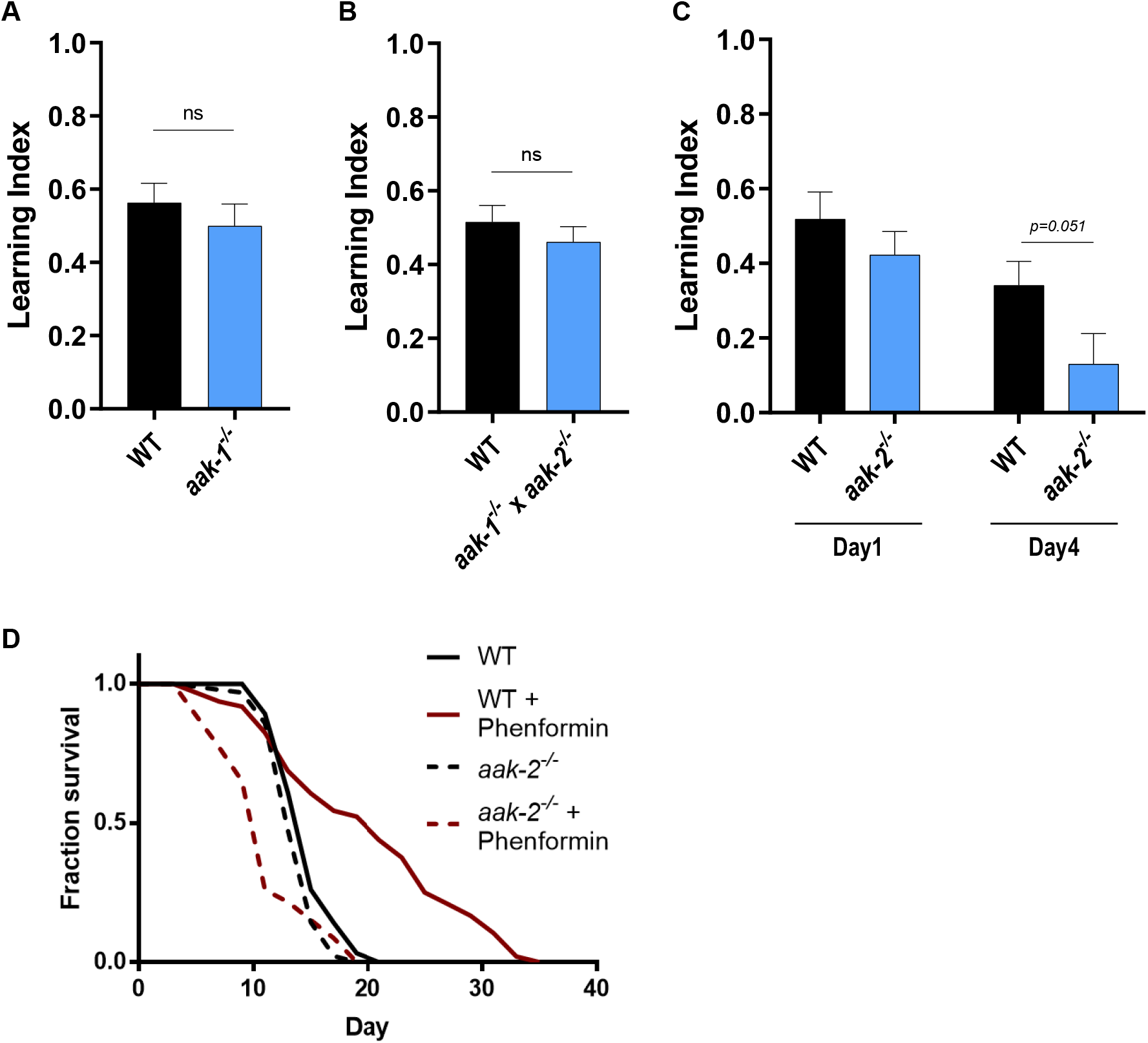
Loss of functional of AMPK does not impair associative learning in young *C. elegans*. **A.** Learning index of associative learning assay comparing day 1 adult WT *C. elegans* and *aak-1^−/−^* mutants. Means WT 0.56 ± 0.05 (n=18), *aak-1^−/−^* 0.50 ± 0.07 (n=17). Comparison WT vs *aak-1^−/−^ p=0.436* **B.** Learning index of associative learning assay comparing day 1 adult WT *C. elegans* and *aak-1^−/−^* × *aak-2^−/−^* mutants. Means WT 0.52 ± 0.04 (n=19), *aak-1^−/−^* × *aak-2^−/−^* 0.46 ± 0.07 (n=19). Comparison WT vs *aak-1^−/−^* × *aak-2^−/−^ p=0.375* **C.** Learning index of associative learning assay comparing day 1 and day 4 adult WT *C. elegans* and *aak-2^−/−^* mutants. Means WT day 1 0.52 ± 0.07 (n=14), *aak-2^−/−^* day 1 0.42 ± 0.06 0. (n=15), WT day 4 0.34 ± 0.06 (n=15), *aak-2^−/−^* day 4 0.13 ± 0.08 (n=15). Comparison WT day 4 vs *aak-2^−/−^* day 4 *p=0.051* **D.** Lifespan assay of WT *C. elegans* and *aak-2^−/−^* mutants, untreated or treated with Phenformin 4.5mM from L4 stage. 2 biological replicates, statistical analysis of all lifespan experiments in SupTable2.

**Supplemental Figure 3:**
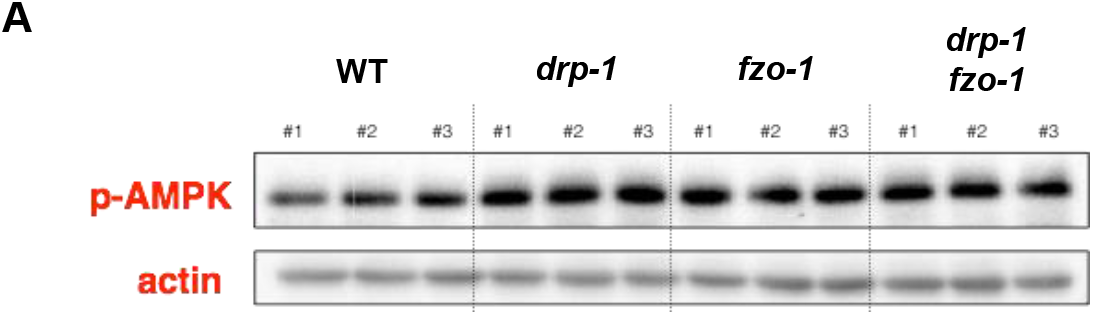
p-AMPK levels in mitochondrial mutant strains. **A.** Western blot of day 1 adult WT *C. elegans*, *drp-1^−/−^* mutants, *fzo-1^−/−^* mutants and *drp-1^−/−^* x *fzo-1^−/−^* mutants. 3 biological replicates were run in parallel. p-AMPK levels were compared and normalized to actin levels.

**Supplemental Figure 4:**
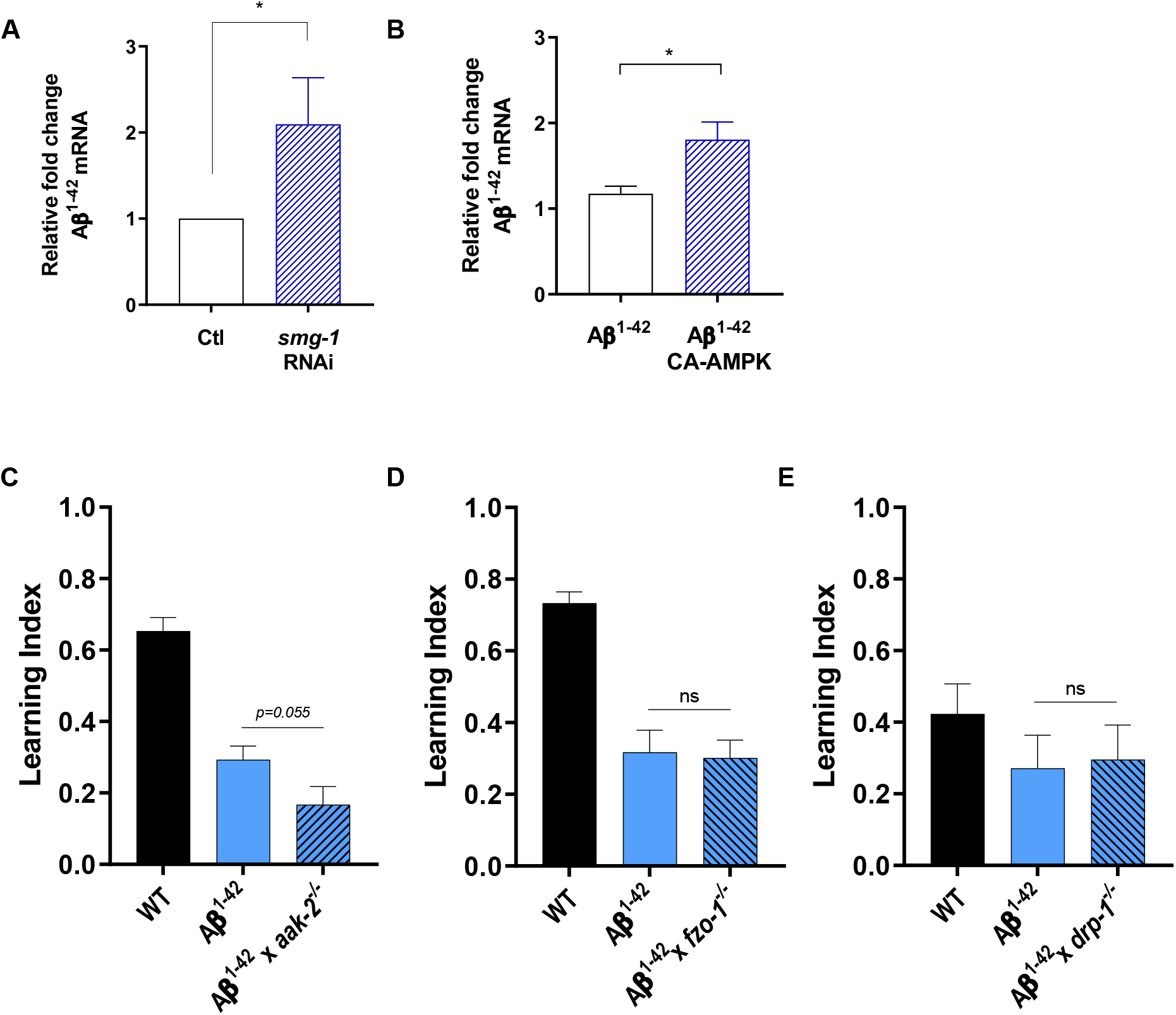
Modulation of mitochondrial networks is not sufficient to alleviate Aβ^1-42^ induced loss of associative learning. **A.** qPCR analysis of day 1 adult Abeta mRNA in Aβ^1-42^ strain grown from hatch on HT115-EV (Ctl) or HT115 expressing *smg-1* RNAi (*smg-1* RNAi). *cdc-42* expression is used as internal control. 3 independent biological replicates. Means Ctl 1, *smg-1* RNAi 2.10 ± 0.31, *p=0.0248* **B.** qPCR analysis of day 1 adult Abeta mRNA in Aβ^1-42^ strain vs CA-AMPK x Aβ^1-42^ strain. *cdc-42* expression is used as internal control. 3 independent biological replicates. Means Aβ^1-42^ 1.17 ± 0.09, CA-AMPK x Aβ^1-42^ 1.81 ± 0.20, *p=0.0465* **C.** Learning index of associative learning assay in day 1 adult WT *C. elegans*, Aβ^1-42^ strain and *aak-2^−/−^* x Aβ^1-42^ strain. Means WT 0.65 ± 0.04 (n=20), Aβ^1-42^ strain 0.29 ± 0.04 (n=20), *aak-2^−/−^* x Aβ^1-42^ strain 0.17 ± 0.05 (n=20). Comparison Aβ^1-42^ strain vs *aak-2^−/−^* x Aβ^1-42^ strain *p=0.055* **D.** Learning index of associative learning assay in day 1 adult WT *C. elegans*, Aβ^1-42^ strain and *fzo-1^−/−^* x Aβ^1-42^ strain. Means WT 0.73 ± 0.03 (n=15), Aβ^1-42^ strain 0.32 ± 0.06 (n=15), *fzo-1^−/−^* x Aβ^1-42^ strain 0.30 ± 0.05 (n=15). Comparison Aβ^1-42^ strain vs *fzo-1^−/−^* x Aβ^1-42^ strain *p=0.844* **E.** Learning index of associative learning assay in day 1 adult WT *C. elegans*, Aβ^1-42^ strain and *drp-1^−/−^* x Aβ^1-42^ strain. Means WT 0.42 ± 0.08 (n=8), Aβ^1-42^ strain 0.27 ± 0.09 (n=9), *drp-1^−/−^* x Aβ^1-42^ strain 0.30 ± 0.09 (n=9). Comparison Aβ^1-42^ strain vs *drp-1^−/−^* x Aβ^1-42^ strain *p=0.857*

**Supplemental Figure 5:**
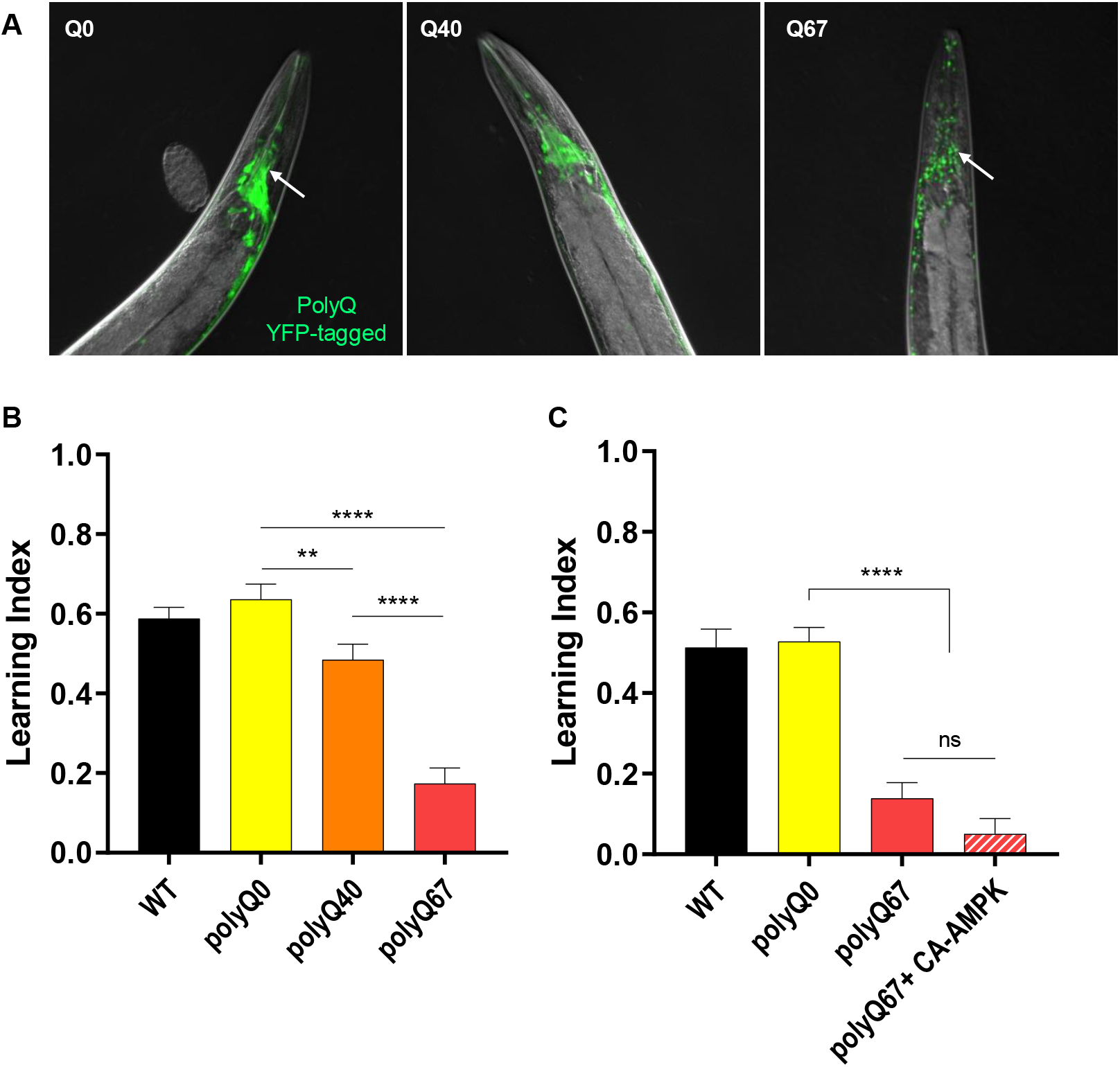
CA-AMPK does not rescue polyQ induced loss of associative learning. **A.** Representative images of polyQ strains (polyQ0, polyQ40, polyQ67) showing CAG repeats dependent aggregation in day 1 *C. elegans* fed on OP50-1 *E. Coli.* Arrows point to either diffuse GFP (left image) or aggregated GFP signal (right image) **B.** Learning index of associative learning assay comparing day 1 adults WT vs strains expressing polyQ0/40/67 under pan neuronal promoter (F25B3.3 promoter). Means WT 0.50 ± 0.03 (n=22), polyQ0 0.64 ± 0.04 (n=25), polyQ40 0.48 ± 0.04 (n=25), polyQ67 0.17 ± 0.04 (n=24). Comparisons polyQ0 vs polyQ40 *p0.075*, polyQ0 vs polyQ67 *p<0.0001*, polyQ40 vs polyQ67 *p<0.001* **C.** Learning index of associative learning assay comparing day 1 adults WT vs polyQ0 vs polyQ67 vs polyQ67 x CA-AMPK. Means WT 0.51 ± 0.05 (n=18), polyQ0 0.53 ± 0.03 (n=20), polyQ67 0.014 ± 0.04 (n=20), polyQ67 x CA-AMPK 0.05 ± 0.04 (n=19). Comparisons polyQ0 vs polyQ67 *p<0.0001*, polyQ0 vs polyQ67 x CA-AMPK *p<0.0001*, polyQ67 vs polyQ67 vs CA-AMPK *p=0.115*.

## ACKNOWLEDGEMENTS

We thank the Caenorhabditis Genetics Center [supported by National Institutes of Health – Office of Research Infrastructure Programs (P40 OD010440)] for strains. We thank Staci Thornton and Juan Ramirez for assistance with learning assays and strain characterization, Gunes Parlakgul for confocal microscopy training and members of the Mair laboratory for scientific discussion and critical reading of the manuscript. We thank Dr. Colleen Murphy and the Murphy Lab at Princeton for training on *C. elegans* behavioral assays.

## MATERIALS AND METHODS

### Worm Strains and Culture

Worms were routinely grown and maintained on standard nematode growth media (NGM) seeded with *E. coli* (OP50-1). *E. coli* bacteria were cultured overnight in LB at 37 °C, after which 100μl of liquid culture was seeded on plates to grow for 2 days at room temperature(81). See Supplemental Table 1 for the complete list of strains generated and employed in this study. As indicated in Supplemental Table 1 some *C. elegans* strains were obtained from the *Caenorhabditis* Genetic Center which is funded by NIH Office of Research Infrastructure Programs (P40 OD010440).

### Lifespan experiments

Lifespan experiments were performed on standard nematode growth media (NGM) at 20°C. Worms were synchronized by timed egg lays using gravid adults or by bleaching gravid adults. When the progeny reached adulthood (72hr), 100 worms (unless otherwise noted) were transferred to fresh plates at 20-25 worms per plate and this was considered time T=0. Worms were transferred to fresh bacterial lawns every other day until the first deaths (10–14 days). Survival was scored every 2 days and a worm was deemed dead when unresponsive to 3 taps on the head and tail. Worms were censored due to contamination on the plate, leaving the NGM, eggs hatching inside the adult or loss of vulval integrity during reproduction. Only in the Phenformin treatment lifespans, 5-Fluoro-2’-deoxyuridine (FUDR) was added to media to prevent excessive censoring. FUDR (100μl; 1 mg/ml) was added on top of the bacterial lawn 24hr before picking worms to the plate at L4 stage, and worms were transferred off FUDR-containing plates once reproduction had ceased (9 days), after which the assays continued normally. Phenformin hydrochloride powder was added in the NGM at a final concentration of 4.5mM. All lifespan repeats are available in supplemental Table 2.

### Western Blots

For each biological sample, approximately 300-500 day 1 adult (unless otherwise noted) worms were collected in M9 buffer with 0.01% Tween-20, washed three times in M9 and snap-frozen in liquid nitrogen. RIPA buffer supplemented with protease inhibitors (Sigma, MI, USA #8340) and phosphatase inhibitors (Roche, Basel, Switzerland #4906845001) was added before lysing the worms via sonication (Qsonica, CT, USA Q700). Lysate was centrifuged at 14,000g for 5min at 4°C and protein concentration was measured using Pierce BCA protein assay kit (Thermo Fisher Scientific, MA, USA PI23227) following manufacturer’s instructions. To denature the proteins, 5X RSB was added and samples were heated to 95°C for 5 min. 20μg of total protein was resolved by SDS–PAGE using 7.5% acrylamide gels and transferred to nitrocellulose membranes. Membranes were blocked in 5% BSA in TBS-T before incubation with primary antibodies: phospho-AMPKα Thr172 (1:1000, Cell signaling, MA, USA #2535), β-actin (1:1000, Abcam, Cambridge, UK #8226), GFP (1:1000, Clontech/Takara, #632380). Secondary antibodies were HRP-conjugated and bands were visualized by the ECL Western Blotting Detection Reagent (GE Healthcare, IL, USA Catalog number: 95038–560) and bands were quantified with Gel Doc system (Bio Rad) and Image Lab software (Version 4.1).

### RNAi

Feeding RNAi clones were obtained from the Ahringer or Vidal RNAi libraries and sequence-verified before using. To use, bacteria were grown overnight in LB broth with 100 μg/mL carbenicillin and 12.5 μg/mL tetracycline, seeded on NGM plates with 100 μg/mL carbenicillin and allowed 48 hours to grow at room temperature. At least 4 hours before use, 0.1M IPTG solution with 100 μg/mL carbenicillin and 12.5 μg/mL tetracycline was added to the bacterial lawn to induce dsRNA expression.

### RNA extraction

For each biological sample, approximately 300-500 day 1 adult (unless otherwise noted) worms were collected in M9 buffer with 0.01% Tween-20, washed three times in M9 and snap-frozen in 600μL of Qiazol (Qiagen, Venlo, Netherlands, #79306) in liquid nitrogen. 3 biological replicates (same bleach) were collected and the experiment was repeated twice. All samples were stored in −80°C freezer until RNA extraction. In order to crack open the cuticle, the samples were thawed in a 37°C water bath for 2 min, vortexed for 30 sec and frozen in liquid nitrogen for 1 min. This cycle was repeated a minimum of 5 times to ensure successful RNA extraction. RNA extraction was performed using QIAGEN RNeasy Mini Kit (QIAGEN, #74104) following manufacturer’s instructions and eluted in 30μL RNAse-free water. RNA concentration was determined by diluting 2μL of RNA in 10mM Tris buffer pH7.5 (1:5 dilution) and using NanoDrop spectrophotometer. Purified RNA samples were stored at −80°C until analysis.

### Quantitative Real-Time PCR

cDNA was synthesized from 200μg of RNA with SuperScript VILO Master Mix (ThermoFisher Scientific, 11755050) following manufacturer’s instructions. 2μL of cDNA (diluted 1:5 in H_2_O) was used as template for each RT-PCR reaction. Three independent biological replicates were used for each genotype/condition, run in technical triplicates and always run in parallel with Taqman control probe to invariant control gene *cdc-42* (Taqman QPCR #4448484) for normalization on a 96 well plate. RT-PCR was performed on the StepOne Plus qPCR Machine (Life Technologies, MA, USA) using Taqman Universal Master Mix II (Life Technologies, 4440040). Taqman probes used to target each gene of interest are as follows: *cdc-42* (Taqman QPCR #4448484), Abeta primers (forward CACCATGAGTCCAATGATTGCA, reverse CACCATGAGTCCAATGATTGCA). Relative expression levels were calculated using ΔΔCt method.

### Genotyping of deletion alleles

Worms were individually lysed in single worm lysis buffer and lysates were used as templates for PCR reactions with a combination of 2–3 primers that will produce bands of different sizes for wild type and mutant alleles. Primers and PCR conditions for each deletion allele are listed in supplemental Table 1.

### Amyloid peptide staining

We used a modified version of the Finney and Ruvkun immunofluorescence protocol (Finney and Ruvkun, Cell, 1990) described by Bettinger et al (Bettinger, Development, 1996). Quickly, to be prepared for the immunohistochemistry, worms were collected in a modified Ruvkun Finney Buffer (RFB: 160 mM KCl, 40 mM NaCl, 20 mM EGTA, 10 mM spermidine, 30 mM PIPES, 50% Methanol) supplemented with 2% paraformaldehyde and snap-froze in a dry-ice and ethanol bath. Samples were then incubated on ice for 3.5h before being treated 4h at 37°C with 1% beta-mercaptoethanol in Tris-Triton Buffer (TTB), then 15min at 37°C with 10mM DTT in BO3 Buffer (0.01 M H3BO3 pH 9.2, 0.01 M NaOH) and finally 15min at room temperature with 0.3% Hydrogen peroxide in BO3 Buffer.

For the immunohistochemistry, the samples were then blocked 30 min at room temperature (Blocking buffer: 0.1% Bovine Serum Albumin, 0.5% Triton-X, 1 mM EDTA, 0.05% Sodium Azide in PBS) and incubate overnight at 4°C with the monoclonal anti-amyloid beta antibody 6E10 (Biolegend, 803001, lot. B228658) at a concentration of 3μg/ml (Antibody buffer: 1% Bovine Serum Albumin, 0.5% Triton-X, 1 mM EDTA, 0.05% Sodium Azide in PBS). The worms were then gently washed in blocking buffer and incubated again overnight with the secondary antibody (CY3-conjugated Goat anti-mouse igG1 specific antibody, 115-165-205, Jackson ImmunoResearch) diluted 1/350 in the antibody buffer. Nuclei were then stained using DAPI and worms were mounted on Superfrost slides with fluoromount mounting medium to be imaged. An average of 100 day 1 nematodes were stained using this technique and 20 nematodes were imaged.

### Drug treatment

In order to treat nematodes with phenformin, we added phenformin hydrochloride (Sigma-Aldrich, #P7045-10G) diluted in distilled water in the standard NGM plate preparation. Stock solution of 200mM diluted in distilled water was prepared and aliquoted in eppendorf tubes. Phenformin solution was kept at −20°C and thawed at RT when needed. Stocks can be kept for 1 month in −20°C. For lifespan assays 10mL plates were used, for associative learning assays 25mL plates were poured with a final phenformin concentration of 4.5mM. In order to treat nematodes with cycloheximide we made a stock solution (1mg/mL) in distilled water and was put on a 10mg NGM plate for a final concentration of 800μg/mL. In order to treat nematodes with actinomycin D, we made a stock solution (1mg/mL) in DMSO and further diluted at 1μg/mL in M9 buffer.

### Microscopy

For imaging of mitochondria in neurons, images were taken in the Sabri Ulker imaging lab using a Yokogawa CSU-X1 spinning disk confocal system (Andor Technology, South Windsor, CT, USA) combined with a Nikon Ti-E inverted microscope (Nikon Instruments, Melville, NY, USA). Images were taken using a 100x/1.45 oil Plan Apo objective lens, Zyla cMOS (Zyla 4.2 Plus USB3) camera and 488 nm Laser for GFP using the same exposure time (70ms). Optical slice thickness was 0.2 μm using the PIEZO technology. NIS elements software was used for acquisition parameters, shutters, filter positions and focus control. An average of 10 nematodes were imaged per condition and per time point.

For imaging of the amyloid peptide staining, images were taken using the AxioImager microscope with an AxioCamMR3 Camera and a 40x/0.75 dry Plan Neofluar objective lens. Exposure time was fixed for all samples: CY3 250ms and DAPI 20ms.

### Statistical analysis

All experiments were analyzed using Graphpad Prism 8 on Windows. For lifespans assays, *p* values were calculated using the Log-rank (Mantel-Cox) statistical test. Unless otherwise noted in the figure legends, data were plotted as Mean +/− SEM. Statistical analysis was performed using unpaired two tail t-test and *p* values were defined as follows: ns no significance, *<0.05, **<0.001, ***<0.001, ****<0.0001

### Learning assay pairing NaCl and starvation

#### Chemotaxis assay

Chemotaxis assays were based on those described by Bargmann and colleagues (82,83) with some modifications. For assay plates composition see below. For NaCl chemotaxis assays, an agar plug was excised from a “high NaCl” plate (see composition below) with the larger end of a 1000μL plastic pipet tip and placed on the surface, on the left side, of an assay plate (containing no NaCl) which was then left for 4hrs at RT. Shortly before the chemotaxis assay, the NaCl plug was removed and 1μL of 1M sodium azide was spotted onto the same position to anaesthetize the animals at the centre of the gradient. As a control, sodium azide was also spotted at the opposite side of the plate (right side, Fig1A). After 4h conditioning, worms were washed off the late with M9 buffer and quickly washed 3 times with M9 buffer before approximately 200 animals (from each condition) were then placed equidistant from these two spots (Fig1A) and left to move freely on the assay plate for 1h at 20 °C. Excess buffer was absorbed with a Kimwipe. The chemotaxis index (CI) was calculated as CI=(N+ - N−)/total number of worms, where N+ is the number of animals within 1cm of the centre of the NaCl gradient and N− is the number within 1cm of the control spot. In each assay, each condition was performed in 6 replicates (6 chemotaxis plates for [no E. coli/0mM NaCl] worms and 6 chemotaxis plates for [no E. coli/50mM NaCl] worms, for each strain studied). The numbers of independent experiments performed are indicated in the figure legends. In each assay, for each strain, the 6 replicates with conditioned worms [no E. coli/50mM NaCl] were normalized to the mean of the control worms [no E. coli/0mM NaCl] in order to calculate the learning index. Each graph is plotted with the N2 learning index defined as 1.

#### Conditioning procedure

Conditioning protocols were based on those described by Saeki et al.(31). The procedure for the learning assay was as follows. Worms were synchronized by bleaching gravid adults fed at for at least 2 generations. Worms were grown on NGM plates seeded with OP50-1 bacteria until day1 adults. The animals were washed off the plates with M9 buffer and transferred to a falcon tube. The animals were washed three times by allowing the adults to fall through the M9 buffer by gravity and replacing the supernatant, including floating small larvae, with fresh M9 buffer. Washing the worms 3 times ensured that no remaining bacteria was transferred to the conditioning plates. For conditioning, washed animals were placed on a conditioning plate (as mentioned in figure legends), and excess fluid was absorbed with a Kimwipe (see composition of plates below). The standard conditions used were: control condition [no E. coli/0mM NaCl] which allowed us to control for the effect of starvation alone and [no E. coli/50mM NaCl]. After incubation at 20 °C for 4 h, the animals were collected again with M9 buffer and chemotaxis was assayed as described above.

#### Composition of the plates

No NaCl conditioning plates and assay plates: 5 mmol/L potassium phosphate pH 6.0, 1 mmol/L CaCl2, 1 mmol/L MgSO4, 20 g/L Agar

NaCl conditioning plate and normal growth plates: 5 mmol/L potassium phosphate pH 6.0, 1 mmol/L CaCl2, 1 mmol/L MgSO4, 20 g/L Agar, 50mmol/L NaCl

High NaCl plate for agar plug: 5 mmol/L potassium phosphate pH 6.0, 1 mmol/L CaCl2, 1 mmol/L MgSO4, 20 g/L Agar, 100 mmol/L NaCl

#### Preparation of aged C. elegans populations

Worm populations were synchronized by bleaching gravid adults and transferring the eggs to 10cm standard NGM plates seeded with OP50-1 bacteria. When worms reached L4 stage (48h), they were transferred to NGM plates supplemented with 5-Fluoro-2’-deoxyuridine (FUDR) for a final concentration of 8.3μg/mL. Plates were seeded with OP50-1 bacteria grown in LB overnight at 37 °C and centrifuged at 4000rpm. Bacterial pellet obtained from 150mL of culture was resuspended in 50μl M9 buffer and seeded on a 10cm FUDR supplemented NGM plate and left to dry overnight. Worms were left on these plates until they reached desired age. Any plate where worms starved due to lack of bacteria was discarded.

## REFERENCES

1. Escoubas CC, Silva-Garcia CG, Mair WB. Deregulation of CRTCs in Aging and Age-Related Disease Risk. Trends Genet. n.d.;33(5):303–21.

2. Moreno-Trevino MG, Castillo-Lopez J, Meester I. Moving away from amyloid Beta to move on in Alzheimer research. Frontiers in aging neuroscience. 2015;7:2.

3. Jan AT, Azam M, Rahman S, Almigeiti AMS, Choi DH, Lee EJ, et al. Perspective Insights into Disease Progression, Diagnostics, and Therapeutic Approaches in Alzheimer’s Disease: A Judicious Update. Front Aging Neurosci. 2017;9:356.

4. Geerts H, Roberts P, Spiros A, Carr R. A strategy for developing new treatment paradigms for neuropsychiatric and neurocognitive symptoms in Alzheimer’s disease. Front Pharmacol. 2013;4:47.

5. Hardy J, Selkoe DJ. The Amyloid Hypothesis of Alzheimer’s Disease: Progress and Problems on the Road to Therapeutics. Science. 2002;297(5580):353–6.

6. Hardy J, Higgins G. Alzheimer’s disease: the amyloid cascade hypothesis. Science. 1992;256(5054):184–5.

7. Selkoe DJ. Soluble oligomers of the amyloid β-protein impair synaptic plasticity and behavior. Behav Brain Res. 2008;192(1):106–13.

8. Braak H, Tredici KD. Alzheimer’s pathogenesis: is there neuron-to-neuron propagation? Acta Neuropathol. 2011;121(5):589–95.

9. Cummings JL, Ringman J, Vinters HV. Neuropathologic correlates of trial-related instruments for Alzheimer’s disease. Am J Neurodegener Dis. 2014;3(1):45–9.

10. Procaccini C, Santopaolo M, Faicchia D, Colamatteo A, Formisano L, Candia P de, et al. Role of metabolism in neurodegenerative disorders. Metabolism: clinical and experimental. n.d.;65(9):1376–90.

11. Jha SK, Jha NK, Kumar D, Ambasta RK, Kumar P. Linking mitochondrial dysfunction, metabolic syndrome and stress signaling in Neurodegeneration. Biochimica et biophysica acta. 21AD;1863(5):1132–46.

12. Wu P, Shen Q, Dong S, Xu Z, Tsien JZ, Hu Y. Calorie restriction ameliorates neurodegenerative phenotypes in forebrain-specific presenilin-1 and presenilin-2 double knockout mice. Neurobiology of aging. n.d.;29(10):1502–11.

13. Witte AV, Fobker M, Gellner R, Knecht S, Floel A. Caloric restriction improves memory in elderly humans. Proceedings of the National Academy of Sciences of the United States of America. 27AD;106(4):1255–60.

14. Burkewitz K, Zhang Y, Mair WB. AMPK at the nexus of energetics and aging. Cell metabolism. 1AD;20(1):10–25.

15. Hardie DG, Ross FA, Hawley SA. AMPK: a nutrient and energy sensor that maintains energy homeostasis. Nature reviews Molecular cell biology. 22AD;13(4):251–62.

16. Mair W, Morantte I, Rodrigues AP, Manning G, Montminy M, Shaw RJ, et al. Lifespan extension induced by AMPK and calcineurin is mediated by CRTC-1 and CREB. Nature. 17AD;470(7334):404–8.

17. Stenesen D, Suh JM, Seo J, Yu K, Lee KS, Kim JS, et al. Adenosine nucleotide biosynthesis and AMPK regulate adult life span and mediate the longevity benefit of caloric restriction in flies. Cell metabolism. 8AD;17(1):101–12.

18. Martin-Montalvo A, Mercken EM, Mitchell SJ, Palacios HH, Mote PL, Scheibye-Knudsen M, et al. Metformin improves healthspan and lifespan in mice. Nature communications. 2013;4(1):2192.

19. Salminen A, Kaarniranta K, Kauppinen A. Age-related changes in AMPK activation: Role for AMPK phosphatases and inhibitory phosphorylation by upstream signaling pathways. Ageing research reviews. n.d.;28:15–26.

20. Greco SJ, Sarkar S, Johnston JM, Tezapsidis N. Leptin regulates tau phosphorylation and amyloid through AMPK in neuronal cells. Biochemical and biophysical research communications. 27AD;380(1):98–104.

21. Thornton C, Bright NJ, Sastre M, Muckett PJ, Carling D. AMP-activated protein kinase (AMPK) is a tau kinase, activated in response to amyloid beta-peptide exposure. Biochem J. 15AD;434(3):503–12.

22. Vingtdeux V, Davies P, Dickson DW, Marambaud P. AMPK is abnormally activated in tangle- and pre-tangle-bearing neurons in Alzheimer’s disease and other tauopathies. Acta neuropathologica. n.d.;121(3):337–49.

23. Ju TC, Chen HM, Chen YC, Chang CP, Chang C, Chern Y. AMPK-alpha1 functions downstream of oxidative stress to mediate neuronal atrophy in Huntington’s disease. Biochimica et biophysica acta. n.d.;1842(9):1668–80.

24. Ju TC, Chen HM, Lin JT, Chang CP, Chang WC, Kang JJ, et al. Nuclear translocation of AMPK-alpha1 potentiates striatal neurodegeneration in Huntington’s disease. The Journal of cell biology. 25AD;194(2):209–27.

25. Xu Y, Liu C, Chen S, Ye Y, Guo M, Ren Q, et al. Activation of AMPK and inactivation of Akt result in suppression of mTOR-mediated S6K1 and 4E-BP1 pathways leading to neuronal cell death in in vitro models of Parkinson’s disease. Cellular signalling. n.d.;26(8):1680–9.

26. Ng CH, Guan MS, Koh C, Ouyang X, Yu F, Tan EK, et al. AMP kinase activation mitigates dopaminergic dysfunction and mitochondrial abnormalities in Drosophila models of Parkinson’s disease. The Journal of neuroscience: the official journal of the Society for Neuroscience. 10AD;32(41):14311–7.

27. Vingtdeux V, Giliberto L, Zhao H, Chandakkar P, Wu Q, Simon JE, et al. AMP-activated protein kinase signaling activation by resveratrol modulates amyloid-beta peptide metabolism. The Journal of biological chemistry. 19AD;285(12):9100–13.

28. Burkewitz K, Morantte I, Weir HJM, Yeo R, Zhang Y, Huynh FK, et al. Neuronal CRTC-1 Governs Systemic Mitochondrial Metabolism and Lifespan via a Catecholamine Signal. Cell. 2015;160(5):842–55.

29. Toyama EQ, Herzig S, Courchet J, L. JrL T, Loson OC, Hellberg K, et al. Metabolism. AMP-activated protein kinase mediates mitochondrial fission in response to energy stress. Science. 15AD;351(6270):275–81.

30. Weir HJ, Yao P, Huynh FK, Escoubas CC, Goncalves RL, Burkewitz K, et al. Dietary Restriction and AMPK Increase Lifespan via Mitochondrial Network and Peroxisome Remodeling. Cell Metab. 2017;26(6):884–896.e5.

31. Saeki S, Yamamoto M, Iino Y. Plasticity if chemotaxis revealed by paired presentation of a chemoattractant and starvation in the nematode Caenorhabditis elegans. The Journal of Experimental Biology. 2001;

32. Ardiel EL, Rankin CH. An elegant mind: learning and memory in Caenorhabditis elegans. Learning & memory. n.d.;17(4):191–201.

33. Kauffman AL, Ashraf JM, Corces-Zimmerman MR, Landis JN, Murphy CT. Insulin signaling and dietary restriction differentially influence the decline of learning and memory with age. PLoS biology. n.d.;8(5):e1000372.

34. Cabreiro F, Au C, Leung KY, Vergara-Irigaray N, Cocheme HM, Noori T, et al. Metformin retards aging in C. elegans by altering microbial folate and methionine metabolism. Cell. 28AD;153(1):228–39.

35. Koshiba T, Detmer SA, Kaiser JT, Chen H, McCaffery JM, Chan DC. Structural Basis of Mitochondrial Tethering by Mitofusin Complexes. Science. 2004;305(5685):858–62.

36. Chen H, Detmer SA, Ewald AJ, Griffin EE, Fraser SE, Chan DC. Mitofusins Mfn1 and Mfn2 coordinately regulate mitochondrial fusion and are essential for embryonic development. J Cell Biology. 2003;160(2):189–200.

37. Roy M, Reddy PH, Iijima M, Sesaki H. Mitochondrial division and fusion in metabolism. Curr Opin Cell Biol. 2015;33:111–8.

38. Fröhlich C, Grabiger S, Schwefel D, Faelber K, Rosenbaum E, Mears J, et al. Structural insights into oligomerization and mitochondrial remodelling of dynamin 1-like protein. Embo J. 2013;32(9):1280–92.

39. Pareyson D, Saveri P, Sagnelli A, Piscosquito G. Mitochondrial dynamics and inherited peripheral nerve diseases. Neurosci Lett. 2015;596:66–77.

40. Rui Y, Tiwari P, Xie Z, Zheng JQ. Acute Impairment of Mitochondrial Trafficking by β-Amyloid Peptides in Hippocampal Neurons. J Neurosci. 2006;26(41):10480–7.

41. Dosanjh LE, Brown MK, Rao G, Link CD, Luo Y. Behavioral phenotyping of a transgenic Caenorhabditis elegans expressing neuronal amyloid-beta. Journal of Alzheimer’s disease: JAD. 2010;19(2):681–90.

42. McColl G, Roberts BR, Gunn AP, Perez KA, Tew DJ, Masters CL, et al. The Caenorhabditis elegans A beta 1-42 model of Alzheimer disease predominantly expresses A beta 3-42. The Journal of biological chemistry. 21AD;284(34):22697–702.

43. Pratley RE, Salbe AD, Ravussin E, Caviness JN. Higher sedentary energy expenditure in patients with Huntington’s disease. Ann Neurol. 2000;47(1):64–70.

44. Milakovic T, Johnson GVW. Mitochondrial Respiration and ATP Production Are Significantly Impaired in Striatal Cells Expressing Mutant Huntingtin. J Biol Chem. 2005;280(35):30773–82.

45. Morley JF, Brignull HR, Weyers JJ, Morimoto RI. The threshold for polyglutamine-expansion protein aggregation and cellular toxicity is dynamic and influenced by aging in Caenorhabditis elegans. Proceedings of the National Academy of Sciences of the United States of America. 6AD;99(16):10417–22.

46. Brignull HR, Moore FE, Tang SJ, Morimoto RI. Polyglutamine proteins at the pathogenic threshold display neuron-specific aggregation in a pan-neuronal Caenorhabditis elegans model. The Journal of neuroscience: the official journal of the Society for Neuroscience. 19AD;26(29):7597–606.

47. Rubinsztein DC, Leggo J, Coles R, Almqvist E, Biancalana V, Cassiman JJ, et al. Phenotypic characterization of individuals with 30-40 CAG repeats in the Huntington disease (HD) gene reveals HD cases with 36 repeats and apparently normal elderly individuals with 36-39 repeats. Am J Hum Genet. 1996;59(1):16–22.

48. Duyao M, Ambrose C, Myers R, Novelletto A, Persichetti F, Frontali M, et al. Trinucleotide repeat length instability and age of onset in Huntington’s disease. Nat Genet. 1993;4(4):387–92.

49. Altman J, Das GD. Autoradiographic and histological evidence of postnatal hippocampal neurogenesis in rats. J Comp Neurol. 1965;124(3):319–35.

50. Castilla-Ortega E, Pedraza C, Estivill-Torrús G, Santín LJ. When is adult hippocampal neurogenesis necessary for learning? Evidence from animal research. Rev Neuroscience. 2011;22(3):267–83.

51. Epp JR, Chow C, Galea LAM. Hippocampus-dependent learning influences hippocampal neurogenesis. Front Neurosci-switz. 2013;7:57.

52. Anderson ML, Sisti HM, Curlik DM, Shors TJ. Associative learning increases adult neurogenesis during a critical period. Eur J Neurosci. 2011;33(1):175–81.

53. Sampedro-Piquero P, Moreno-Fernández RD, Mañas-Padilla MC, Gil-Rodríguez S, Gavito AL, Pavón FJ, et al. Training memory without aversion: appetitive hole-board spatial learning increases adult hippocampal neurogenesis. Neurobiol Learn Mem. 2018;151:35–42.

54. Chen X, Barclay JW, Burgoyne RD, Morgan A. Using C. elegans to discover therapeutic compounds for ageing-associated neurodegenerative diseases. Chemistry Central journal. 2015;9(1):65.

55. Poels J, Spasic MR, Callaerts P, Norga KK. Expanding roles for AMP-activated protein kinase in neuronal survival and autophagy. Bioessays. n.d.;31(9):944–52.

56. Ronnett GV, Ramamurthy S, Kleman AM, Landree LE, Aja S. AMPK in the brain: its roles in energy balance and neuroprotection. J Neurochem. n.d.;109 Suppl 1(s1):17–23.

57. Spasić MR, Callaerts P, Norga KK. AMP-Activated Protein Kinase (AMPK) Molecular Crossroad for Metabolic Control and Survival of Neurons. Neurosci. 2009;15(4):309–16.

58. Potter WB, O’Riordan KJ, Barnett D, Osting SM, Wagoner M, Burger C, et al. Metabolic regulation of neuronal plasticity by the energy sensor AMPK. PloS one. 1AD;5(2):e8996.

59. Samuel MA, Voinescu PE, Lilley BN, Cabo R de, Foretz M, Viollet B, et al. LKB1 and AMPK regulate synaptic remodeling in old age. Nature neuroscience. n.d.;17(9):1190–7.

60. Kim J, Park YJ, Jang Y, Kwon YH. AMPK activation inhibits apoptosis and tau hyperphosphorylation mediated by palmitate in SH-SY5Y cells. Brain research. 18AD;1418:42–51.

61. Domise M, Didier S, Marinangeli C, Zhao H, Chandakkar P, Buee L, et al. AMP-activated protein kinase modulates tau phosphorylation and tau pathology in vivo. Scientific reports. 27AD;6(1):26758.

62. Ma T, Chen Y, Vingtdeux V, Zhao H, Viollet B, Marambaud P, et al. Inhibition of AMP-activated protein kinase signaling alleviates impairments in hippocampal synaptic plasticity induced by amyloid beta. The Journal of neuroscience: the official journal of the Society for Neuroscience. 3AD;34(36):12230–8.

63. DiTacchio KA, Heinemann SF, Dziewczapolski G. Metformin treatment alters memory function in a mouse model of Alzheimer’s disease. Journal of Alzheimer’s disease: JAD. 2015;44(1):43–8.

64. Du LL, Chai DM, Zhao LN, Li XH, Zhang FC, Zhang HB, et al. AMPK activation ameliorates Alzheimer’s disease-like pathology and spatial memory impairment in a streptozotocin-induced Alzheimer’s disease model in rats. Journal of Alzheimer’s disease: JAD. 2015;43(3):775–84.

65. Wang Z, Wilson WA, Fujino MA, Roach PJ. Antagonistic Controls of Autophagy and Glycogen Accumulation by Snf1p, the Yeast Homolog of AMP-Activated Protein Kinase, and the Cyclin-Dependent Kinase Pho85p. Mol Cell Biol. 2001;21(17):5742–52.

66. Nixon RA. Autophagy, amyloidogenesis and Alzheimer disease. J Cell Sci. 2007;120(23):4081–91.

67. Lee J-H, Yu WH, Kumar A, Lee S, Mohan PS, Peterhoff CM, et al. Lysosomal Proteolysis and Autophagy Require Presenilin 1 and Are Disrupted by Alzheimer-Related PS1 Mutations. Cell. 2010;141(7):1146–58.

68. Pickford F, Masliah E, Britschgi M, Lucin K, Narasimhan R, Jaeger PA, et al. The autophagy-related protein beclin 1 shows reduced expression in early Alzheimer disease and regulates amyloid β accumulation in mice. J Clin Invest. 2008;118(6):2190–9.

69. Jaeger PA, Wyss-Coray T. Beclin 1 Complex in Autophagy and Alzheimer Disease. Arch Neurol-chicago. 2010;67(10):1181–4.

70. Moreira PI, Carvalho C, Zhu X, Smith MA, Perry G. Mitochondrial dysfunction is a trigger of Alzheimer’s disease pathophysiology. Biochimica Et Biophysica Acta Bba - Mol Basis Dis. 2010;1802(1):2–10.

71. Sweeney G, Song J. The association between PGC-1α and Alzheimer’s disease. Anat Cell Biology. 2015;49(1):1–6.

72. Dolma K, Iacobucci GJ, Zheng KH, Shandilya J, Toska E, White JA, et al. Presenilin influences glycogen synthase kinase-3 β (GSK-3β) for kinesin-1 and dynein function during axonal transport. Hum Mol Genet. 2014;23(5):1121–33.

73. Pagani L, Eckert A. Amyloid-Beta Interaction with Mitochondria. Int J Alzheimer’s Dis. 2011;2011:925050.

74. Hirai K, Aliev G, Nunomura A, Fujioka H, Russell RL, Atwood CS, et al. Mitochondrial Abnormalities in Alzheimer’s Disease. J Neurosci. 2001;21(9):3017–23.

75. Khan SM, Cassarino DS, Abramova NN, Keeney PM, Borland MK, Trimmer PA, et al. Alzheimer’s disease cybrids replicate β-amyloid abnormalities through cell death pathways. Ann Neurol. 2000;48(2):148–55.

76. Mao P, Reddy PH. Aging and amyloid beta-induced oxidative DNA damage and mitochondrial dysfunction in Alzheimer’s disease: Implications for early intervention and therapeutics. Biochimica Et Biophysica Acta Bba - Mol Basis Dis. 2011;1812(11):1359–70.

77. Wang X, Su B, Lee H, Li X, Perry G, Smith MA, et al. Impaired Balance of Mitochondrial Fission and Fusion in Alzheimer’s Disease. J Neurosci. 2009;29(28):9090–103.

78. Barsoum MJ, Yuan H, Gerencser AA, Liot G, Kushnareva Y, Gräber S, et al. Nitric oxide-induced mitochondrial fission is regulated by dynamin-related GTPases in neurons. Embo J. 2006;25(16):3900–11.

79. Wang X, Su B, Siedlak SL, Moreira PI, Fujioka H, Wang Y, et al. Amyloid-β overproduction causes abnormal mitochondrial dynamics via differential modulation of mitochondrial fission/fusion proteins. Proc National Acad Sci. 2008;105(49):19318–23.

80. Teo E, Ravi S, Barardo D, Kim H-S, Fong S, Cazenave-Gassiot A, et al. Metabolic stress is a primary pathogenic event in transgenic Caenorhabditis elegans expressing pan-neuronal human amyloid beta. Elife. 2019;8:e50069.

81. Brenner S. The Genetics of Caenorhabditis elegans. Genetics. 1974;77:71–94.

82. Bargmann CI, Horvitz HR. Chemosensory neurons with overlapping functions direct chemotaxis to multiple chemicals in C. elegans. Neuron. 1991;7(5):729–42.

83. Bargmann CI. Genetic and Cellular Analysis of Behavior in C. Elegans. Annu Rev Neurosci. 1993;16(1):47–71.

